# Circadian rhythm distinctness predicts academic performance based on large-scale learning management system data

**DOI:** 10.64898/2025.12.13.694121

**Authors:** Patrycja Scislewska, Piotr Bebas, Iwona Szatkowska, Rafal Stryjek, Aaron E. Schirmer

**Author notes:** Corresponding Authors: Patrycja Scislewska,; Aaron E. Schirmer.

## Abstract

The subjective amplitude of circadian oscillations (distinctness) is an understudied dimension of circadian rhythmicity, which describes how strongly mood and cognition fluctuate during the day. Emerging evidence suggests that distinctness may be as important as chronotype in regulating the temporal organization of key physiological processes. Previous studies have relied on questionnaires and lacked an objective measure of distinctness. Here, we propose the first objective behavioral measure of distinctness based on circular statistics. We applied this approach to 3.4 million login events from 13,894 unique university students and found a non-linear association between distinctness and academic performance: students with moderate daily rhythmicity achieved the highest performance. This relationship varied by chronotype: larks benefited from stronger rhythms, finches from moderate rhythms, and owls from weaker, more flexible rhythms. Circadian distinctness was also closely linked to social jetlag, which increased with more rigid rhythmicity across chronotypes. These findings suggest that the academic disadvantage often attributed to specific chronotypes does not stem from time preference itself, but from the overall interplay of chronotype, distinctness, and schedules that are incompatible with individual biological timings. Considering rhythm flexibility alongside chronotype may therefore improve educational design and equity.

## Introduction

Circadian rhythms organize nearly every aspect of our physiology and behavior by synchronizing internal biological oscillations with the 24-hour cycles of light and darkness^1–7^. Chronobiological researchers describe these oscillations using quantitative measures like phase, period, and amplitude^8^. In humans, circadian behavior is traditionally characterized using two core parameters: period and phase. Period describes the length of the internal cycle, while phase reflects its timing relative to the external day-night cycle. In human field studies, period offers limited explanatory power because daily light exposure keeps most individuals tightly entrained to a ∼24-hour cycle, leaving little inter-individual variability in the entrained period^9,10^. As a result, research has focused primarily on phase-based measures, particularly chronotype, which captures whether a person tends to be active earlier or later in the day^11,12^. Yet phase alone cannot fully describe how circadian rhythms manifest in daily life^13,14^. Chronotype captures *when* in the day a person tends to function best, but it does not describe *how strongly* their daily rhythm fluctuates. This necessitates greater attention to an underutilized, but potentially consequential, dimension of circadian rhythm: the subjective amplitude of the rhythm. This complementary dimension, also referred to as “distinctness”, reflects how sharply an individual transitions between high- and low-activity states across the day^14–16^. Early works have examined distinctness almost exclusively through questionnaires^15–19^, suggesting that some people experience pronounced fluctuations in mood, energy or cognitive efficiency, whereas others show relatively stable functioning throughout the day. However, despite growing interest, a universally applicable objective behavioral measure of circadian distinctness has been lacking.

Temporal patterns in human behavior can be analysed using circular statistics, a mathematical approach designed for data that repeat in cycles. Unlike standard linear methods, circular analyses account for the fact that time is cyclical – for example, 1 a.m. and 11 p.m. are close together, not at opposite ends of a scale^20^. By converting clock times into angles on a circle, this framework preserves the natural continuity of the 24-hour cycle. Circular statistical tools are therefore well suited for capturing rhythmicity in behavioral timing^21^. If our interest is in understanding the concentration of circadian activity and how it varies across the days, the mean resultant length (*R̅*) a key circular statistic that measures the concentration or dispersion of directional data, would be an appropriate measure^22^. For this reason, we propose that *R̅* is a promising objective measure of circadian distinctness and we will use "*R̅*" and “distinctness” interchangeably throughout this manuscript.

Circadian traits have been repeatedly linked to educational success. Evening-oriented students (so called “Owls”) typically show lower academic performance than morning-oriented (“Larks”) or moderate students (“Finches”)^5,23–25^ . Moreover, misalignment between internal time and institutional schedules, known as social jetlag (SJL), have been consistently associated with poorer educational outcomes especially amongst evening-oriented students^25–27^. Far less is known about the relationship between distinctness and learning. Emerging evidence suggests that high circadian distinctness may coincide with notable psychological and neural patterns. Distinctness has been shown to be negatively related to conscientiousness^28–30^, which in turn is associated with educational performance^31,32^. Moreover, distinctness correlates with greater neuroticism, avoidance behavior, sensitivity to punishment and negative emotionality^14,33–35^, and may be reflected in both brain structure^36^ and brain activity during reward-punishment processing^35^. These data suggest that students with very pronounced daily activity patterns may struggle with course work during their nonoptimal periods, potentially hindering sustained attention and learning but this relationship has not been directly tested.

To address this gap, we utilized a large behavioral dataset of 3.4 million Learning Management System (LMS) login events generated by 13,894 university students. Because students could access the LMS at any time, the dataset consists of naturally occurring, unmanipulated behavioral records and consequently offers high ecological validity, which has been previously shown to reliably reflect students’ internal circadian preferences^25,37,38^. Our main goals were (I) to identify an objective measure of behavioral distinctness based on the temporal distribution of students’ login patterns and (II) to examine its relationship with academic performance. To quantify daily rhythmicity, we represented login times using circular statistics and summarized their concentration using a mean resultant length (*R̅*) calculation. We then examined how *R̅*, chronotype and social jetlag independently and jointly influence students’ academic performance (measured as grade point averages, or GPA) using a multilevel modelling framework. We hypothesize that circadian distinctness will strongly influence academic performance and that an individual’s circadian distinctness is a crucial factor that should be incorporated into studies at the intersection of chronobiology and education.

## Results

### Dataset characteristics

We assessed the distribution of login activity across the 24-hour day using the Rayleigh, Watson U², and Hodges-Ajne tests. Students who had fewer than 26 logins per term were excluded, based on the thresholds derived from the Monte-Carlo simulations of null distributions of *R̅* values. The full filtering procedure is provided in the Methods section (see *Preprocessing*) and Supplementary Materials S3. The initial dataset comprised 33,329 students and approximately 3.4 million login events across 4 semesters (representing 13,894 unique students). After filtering, 17,376 students across 4 semesters (9014 unique students) remained. Among these, 96.2% showed a unimodal distribution of login times, consistent with expected non-uniform daily activity patterns corresponding to active and inactive (sleep-related) phases during the day. Additional characteristics of the dataset before and after filtering are presented in Supplementary Materials S4.

### Circadian characteristics

To quantify the distinctness of the circadian rhythm, we focused on the mean resultant length (*R̅*), a measure of concentration in circular statistics. Exact formulas for this calculation are provided in Supplementary materials S1, S2. Chronotype classification and SJL definitions followed our previous work^25^.

According to Normality tests Shapiro-Wilk and D’Agostino-Pearson normality tests, *R̅* values significantly deviated from normality (*p* < 0.05). Distributions and details are shown in Supplementary Fig. S5. Kruskal-Wallis tests and pairwise Mann-Whitney U tests followed by FDR multiple comparisons correction, revealed differences between seasons, sex, and chronotype, but effect sizes were very small (ε^2^ = 0.001-0.065). Additional details are provided in Supplementary Materials S7, S8, S9.

### Circadian distinctness and academic performance

To characterise how circadian distinctness relates to academic outcomes, we implemented a structured series of statistical models. Below we report the following relationships: (I) *R̅* and GPA across all students, (II) *R̅* and GPA within each chronotype group (Larks, Finches, Owls), (III) *R̅* and SJL across all students, (IV) *R̅* and SJL within each chronotype group, (V) *R̅* and both, GPA and SJL, across all students, (VI) *R̅* and both GPA and SJL, within each chronotype group. This hierarchical approach enabled us to determine not only the overall effects of circadian distinctness, but also how these effects vary across chronotypes with distinct biological and behavioral profiles.

### ***R̅*** and GPA across all students

The relationship between *R̅* and academic performance (GPA) was assessed using Generalized Estimating Equations (GEE) to account for repeated semesters within students. Across 17,376 individuals (9014 unique students), *R̅* showed a significant inverted-U-shaped relationship with GPA (*β* = −1.456, *p* < 0.001). There was no significant linear relationship. The estimated peak regularity was *R̅* ≈ 0.47, indicating that moderate distinctness is associated with the highest average GPA. We visualized the *R̅*-GPA relationship using the LOESS smoothing separately for each semester, season and sex in Fig. 1. The shape of these relationships was unchanged suggesting that semester, season and sex do not impact the quadratic relationship between GPA and *R̅*. Additional details about these models and results can be found in Supplementary Material S10.

**Fig. 1.**
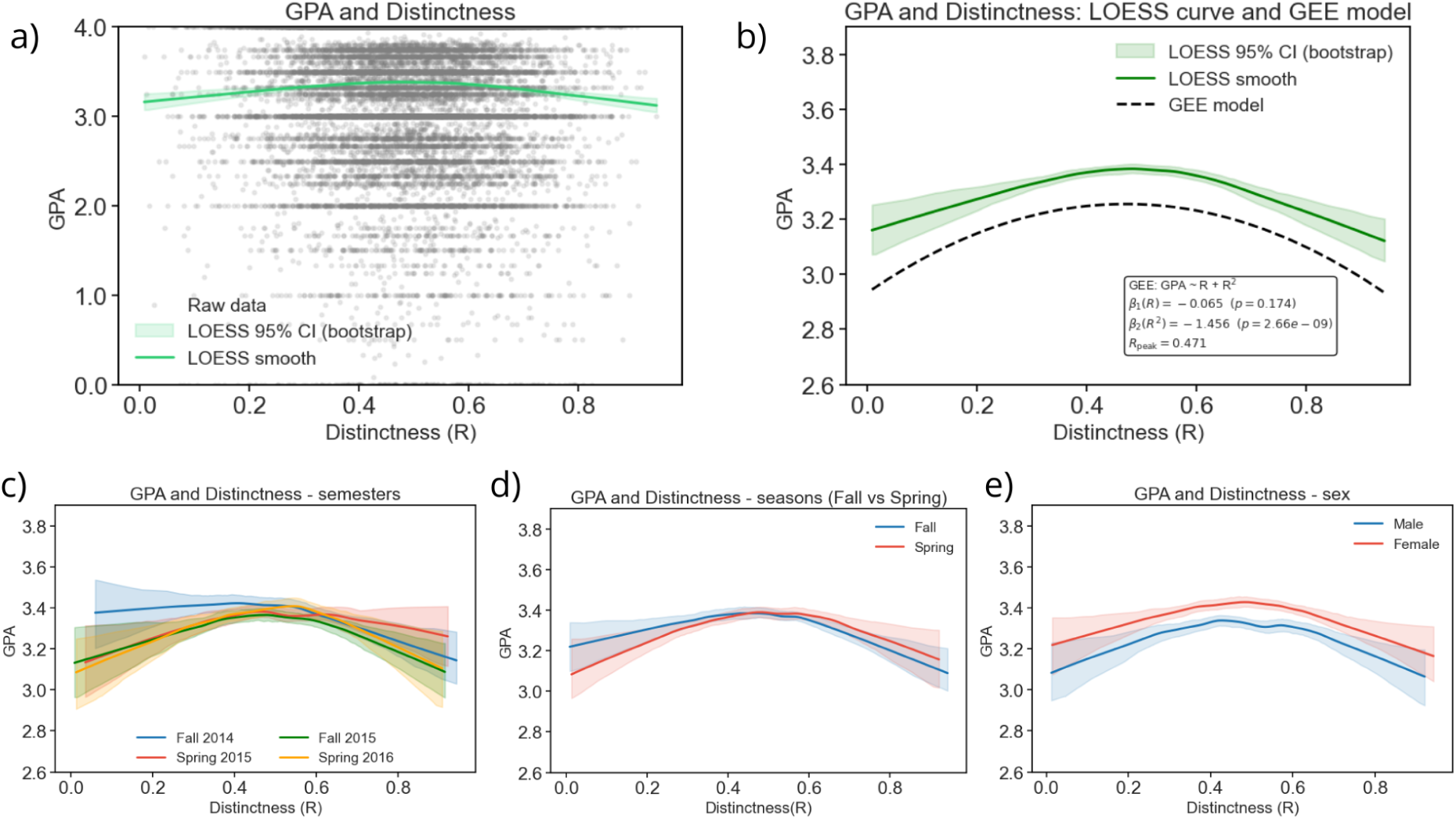
The relationship between *R̅* and GPA. GPA values increase with *R̅* up to about *R̅* = 0.47, after which further increase in circadian distinctness was associated with slightly lower GPA. a) Scatterplot of individual data points (each student from our dataset is marked with a grey dot), overlaid with a LOESS smoothing curve (green solid line) and pointwise 95% bootstrap confidence intervals (1,000 iterations). b) LOESS curve with corresponding 95% confidence bands, shown together with the predicted mean function from the quadratic GEE model (dashed line), statistical estimates for the GEE model are reported in the inset. c-e) LOESS curves with 95% confidence bands calculated separately for semesters, seasons, and sex. Across all subgroup analyses, the shape of the *R̅*–GPA relationship remains consistent, indicating that these factors do not alter the underlying quadratic association between GPA and *R̅*. Abbreviations: R - *R̅*, distinctness of the circadian rhythm, GPA - grade point average

### ***R̅*** and GPA within chronotype groups

Subsequently, we checked if the shape of the *R̅*-GPA relationship differed between chronotypes. We used separate models including linear and quadratic terms for *R̅* for each chronotype group. To determine whether linear or quadratic models fit better within each chronotype group, we compared GEE models using the Quasi-Information Criterion (QIC) ^39^. Results of the QIC are provided in Supplementary table S11.

We found that the *R̅*-GPA relationship differed by chronotype. Among Finches, we observed a clear quadratic relationship (inverted U-shape, *β* = −1.261, *p* < 0.001). In Larks, we found positive linear (*β* = 0.368, *p* < 0.001) and negative quadratic (*β* = −1.064, *p* = 0.016) relationships. In contrast, Owls showed a lower GPA with increasing *R̅*, as indicated by the negative linear trend-level relationship (*β* = –0.333, *p* = 0.059). There was no quadratic relationship in the case of Owls. Together, these results indicate that the impact of circadian rhythm distinctness *R̅* on academic outcomes is chronotype-dependent, with Larks benefiting from higher stability (with optimum *R̅* ∼ 0.6) and Owls showing the opposite pattern - linearly decreasing GPA with increasing *R̅* value (Fig. 2.). Detailed results are shown in Supplementary table S12.

**Fig. 2.**
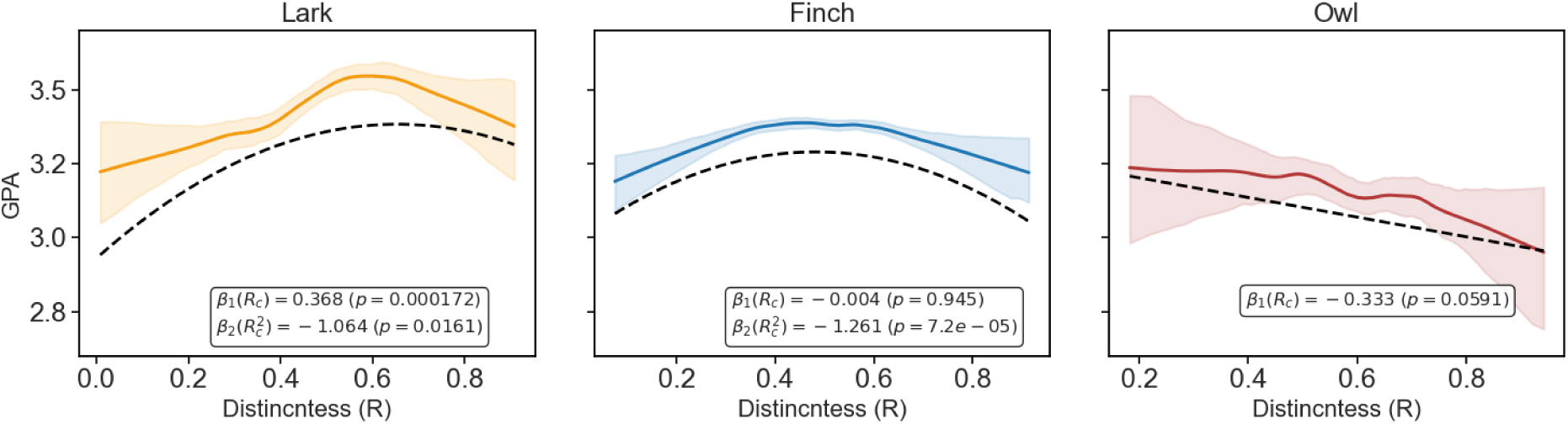
The relationship between *R̅* and GPA within each chronotype group. Each plot shows LOESS curves with corresponding 95% confidence bands, together with the predicted mean function from the GEE model (dashed line), statistical estimates for each GEE model are reported in the inset. Abbreviations: R - *R̅*, distinctness of the circadian rhythm, GPA - grade point average

### ***R̅*** and SJL across all students

Similarly to what we did in the case of *R̅* and GPA, we employed GEE quadratic model to assess the relationship between *R̅* and SJL. We found that SJL increases with an increase in *R̅* (both linear (*β* = 1.134, *p* < 0.001) and quadratic (*β* = −0.847, *p* = 0.002) terms were significant). Again, semester, season (both falls and both springs combined) and sex do not impact the shape of the relationship between SJL and *R̅*. Detailed model results are in Fig. 3. and in Supplementary Materials S13.

**Fig. 3.**
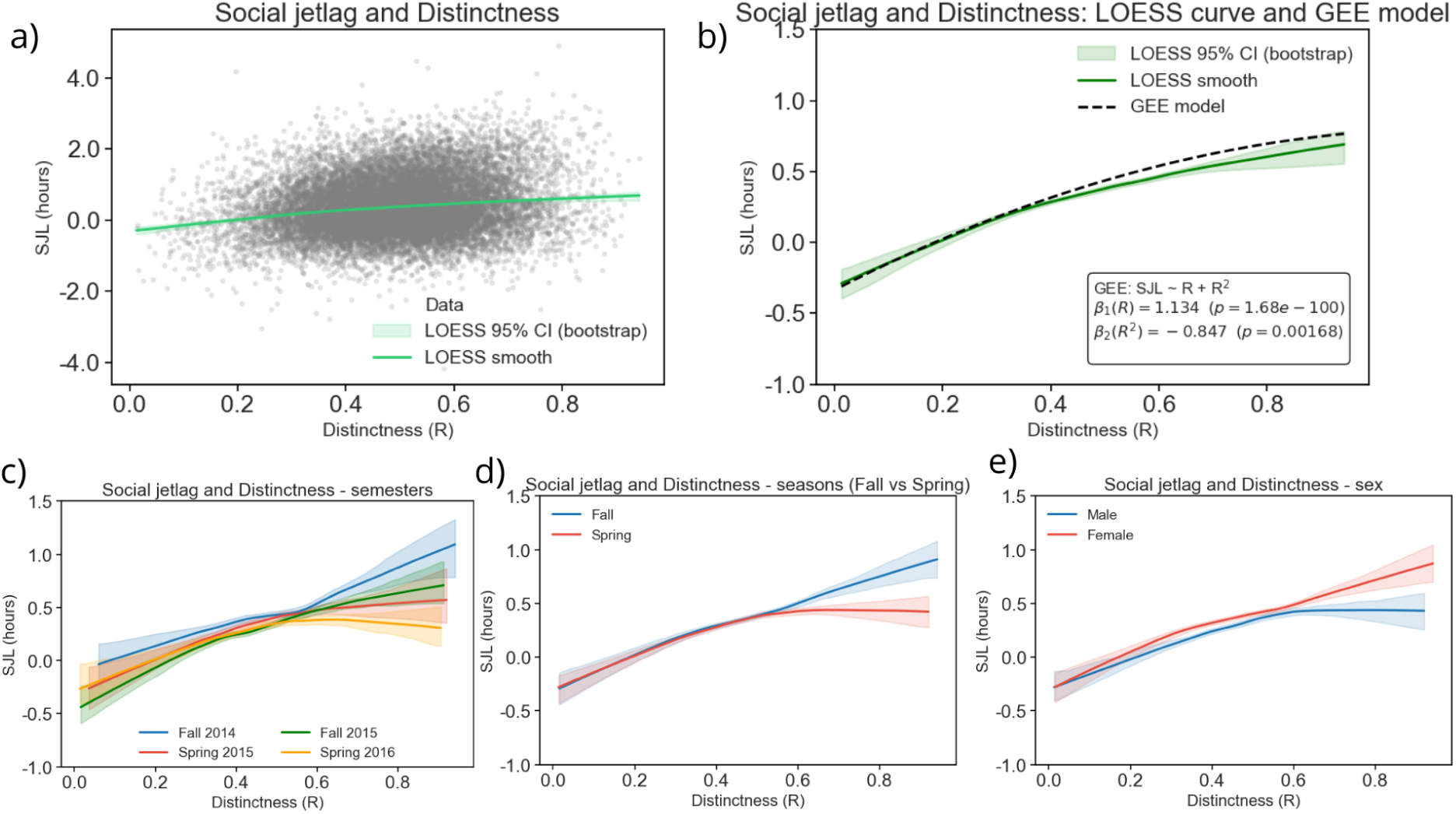
The relationship between *R̅* and SJL. We observed that SJL increases curvilinearly with *R̅*. a) Scatterplot of individual data points (each student from our dataset is marked with a grey dot), overlaid with a LOESS smoothing curve (green solid line) and pointwise 95% bootstrap confidence intervals (1,000 iterations). b) LOESS curve with corresponding 95% confidence bands, shown together with the predicted mean function from the quadratic GEE model (dashed line), statistical estimates for the GEE model are reported in the inset. c-e) LOESS curves with 95% confidence bands calculated separately for semesters, seasons, and sex. Across all subgroup analyses, the shape of the *R̅*–SJL relationship remains consistent, indicating that these factors do not alter the underlying association between SJL and *R̅*. Abbreviations: R - *R̅*, distinctness of the circadian rhythm, SJL - social jetlag.

### ***R̅*** and SJL within chronotype groups

Following our approach from the *R̅*-GPA analysis, we checked if the shape of the *R̅*-SJL relationship also differed between chronotypes. We used separate models including linear and quadratic terms for *R̅* for each chronotype group and compared models using QIC. Results of the QIC are provided in Supplementary table S14.

The *R̅*-SJL relationship differed by chronotype. Among Larks we observed a negative quadratic relationship (*β* = -0.714, *p* = 0.040), among finches – positive linear relationship (*β* = 0.648, *p* < 0.001 in Finches). In contrast, Owls showed a curvilinear increase in SJL (linear *β* = 0.522, *p* = 0.006, quadratic *β* = 4.434, *p* < 0.001). Detailed results are shown in Fig. 4. and Supplementary Table S15. Note that in Fig 4. the y-axis ranges were adjusted separately for each chronotype to improve visualization of the SJL distribution, and owls generally exhibited more than one hour greater SJL compared with larks and finches.

**Fig. 4.**
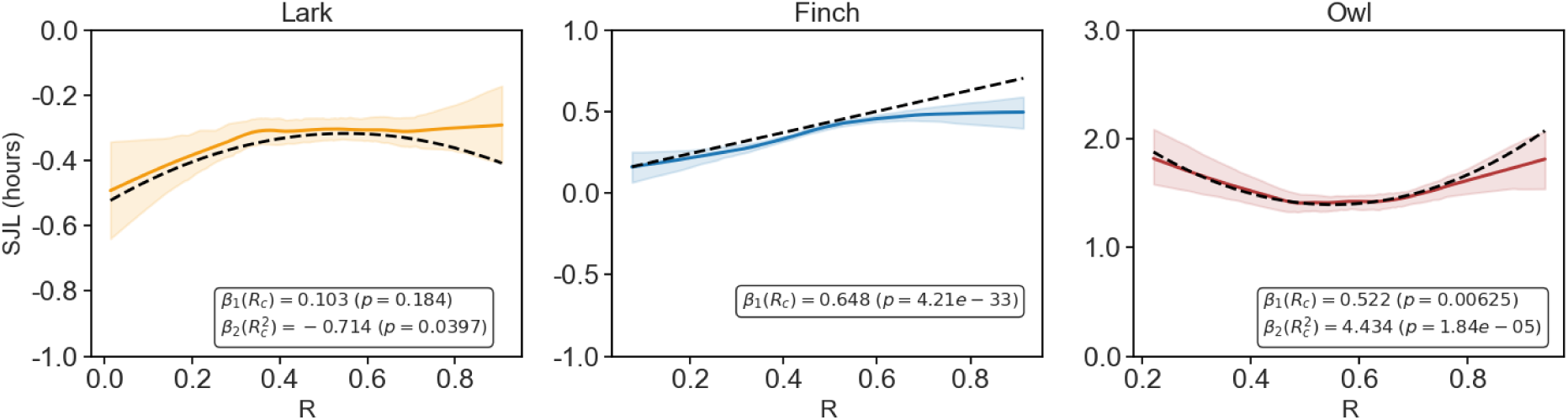
The relationship between *R̅* and SJL within each chronotype group. Each plot shows LOESS curves with corresponding 95% confidence bands, together with the predicted mean function from the GEE model (dashed line), statistical estimates for each GEE model are reported in the inset. Note that the y-axis range was adjusted separately for each chronotype to improve visualization of the SJL distribution and enhance clarity of the plotted relationships.Abbreviations: R - *R̅*, distinctness of the circadian rhythm, SJL - social jetlag.

### ***R̅*** and both, GPA and SJL, across all students

We examined the joint association between *R̅*, SJL and GPA. Both, linear and quadratic effects were significant, indicating nonlinear relationships: GPA declined at extreme levels of both, *R̅* (linear *β* = 1.346, *p* < 0.001, quadratic *β* = -1.303, *p* < 0.001 ) and SJL (linear *β* = –0.077, *p* < 0.001, quadratic β = -0.044, p = 0.011). This indicates that very low or very high level of *R̅* and increased levels (either advance or delayed)of SJL were associated with lower academic performance, whereas moderate values (which here represent little to no SJL) were beneficial, Fig. 5. Additional details are shown in supplementary materials S15.

**Fig. 5.**
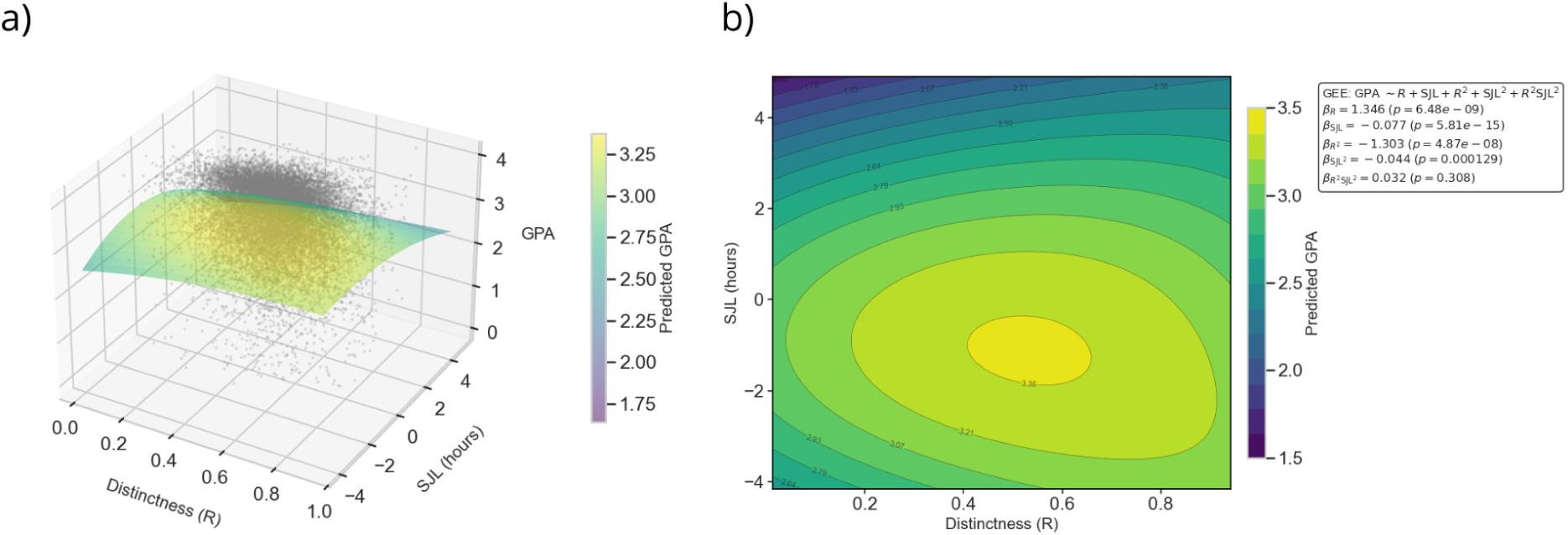
The association of *R̅*, SJL and GPA. GPA varies jointly with *R̅* and SJL, revealing the nonlinear relationships captured by the model. a) Three-dimensional GEE surface illustrating the predicted GPA as a function of *R̅* and SJL. Raw data points are shown in grey, and the semi-transparent surface represents the fitted mean function from the GEE model including linear, quadratic, and interaction terms. b) A contour map of the same GEE surface, showing isolines of predicted GPA across the *R̅*-SJL plane. Warmer colors indicate higher predicted GPA, and contour labels denote estimated GPA values. A statistical summary of the GEE model coefficients is provided in the inset. Abbreviations: R - *R̅*, distinctness of the circadian rhythm, SJL - social jetlag, GPA - grade point average.

### ***R̅*** and both, GPA and SJL, within chronotype groups

Finally, we examined whether the associations between *R̅*, SJL, and GPA differed across chronotype groups. The GEE model revealed several chronotype-specific linear and nonlinear effects.

Finches seem to have a strong association in GPA with *R̅* (linear *β* = 1.427, *p* < 0.001, quadratic *β* = -1.417, *p* < 0.001) and weaker with SJL (linear *β* = -0.081, *p* < 0.001, quadratic *β* = -0.045, *p* = 0.007) meaning that GPA decline at both, extremely high and low, levels of *R̅* and SJL. Larks showed a significant curvilinear association only with *R̅* (linear *β* = 1.547, *p* < 0.001, quadratic *β* = -1.182, p = 0.007) with higher circadian distinctness predicting better academic performance. There was no significant association with SJL. Owls were characterized by two trend-level associations - weak quadratic relationship with SJL (*β* = -0.063, *p* =0.065), and higher-order interaction between the quadratic components of *R̅* and SJL (*β* = 0.102, *p* = 0.09). An interaction between rhythm distinctness and social jetlag emerged only in evening types, indicating a potential chronotype-specific nonlinear relationship, although this effect did not meet conventional significance levels (Fig. 6). Detailed results are provided in Supplementary materials S16.

**Fig. 6.**
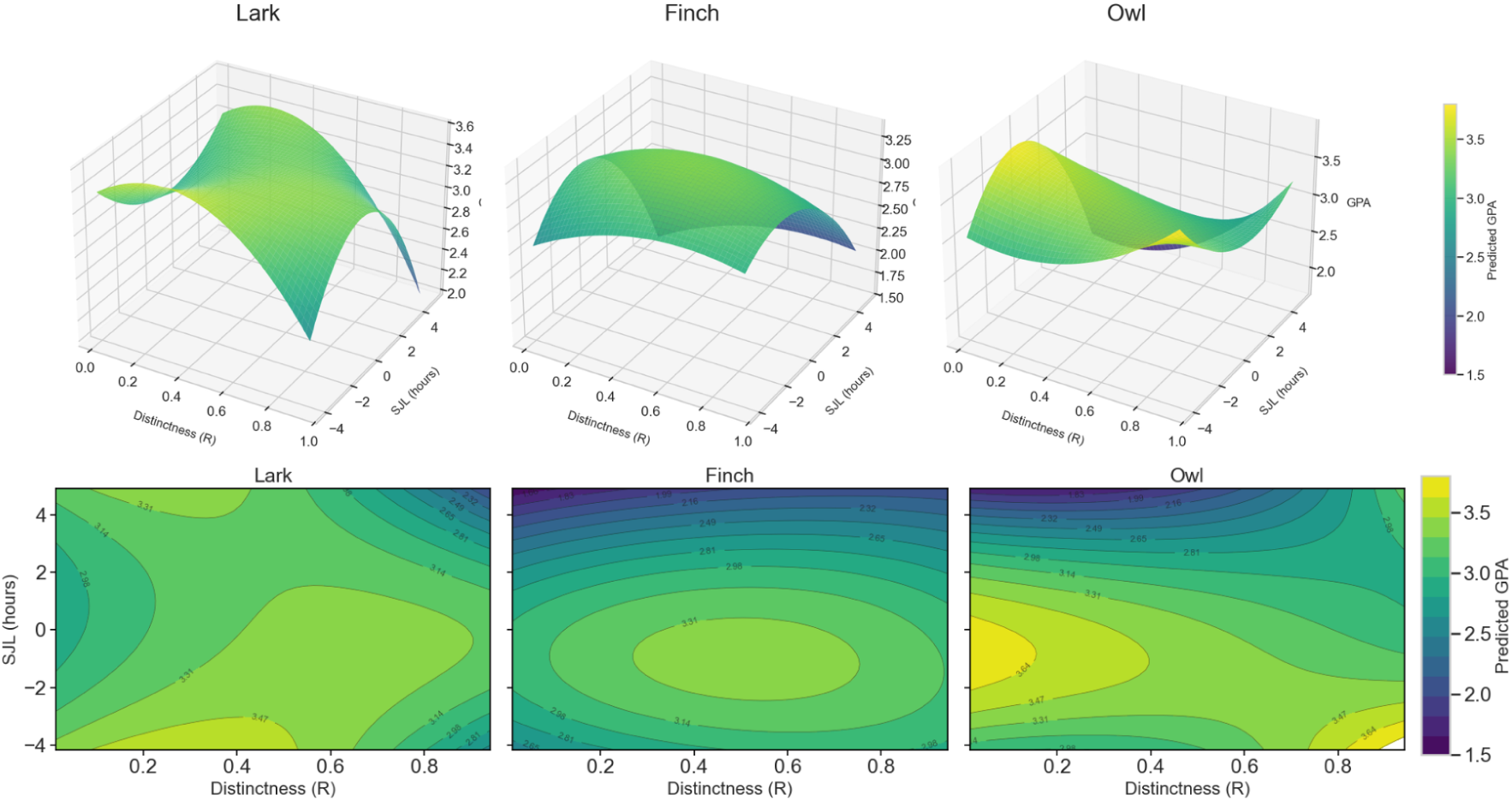
Chronotype-specific predicted GPA surfaces as a function of *R̅* and SJL. a) Three-dimensional GEE surfaces illustrating the predicted GPA as a function of *R̅* and SJL separately for Larks, Finches, and Owls. Each panel shows the fitted mean surface from the chronotype-specific GEE model, with the x-axis representing *R̅*, the y-axis representing SJL (hours), and the z-axis showing predicted GPA. A shared color scale allows direct comparison of the magnitude and shape of the *R̅*–SJL–GPA relationship across chronotypes. b) Corresponding 2D contour maps displaying the same GEE-predicted GPA surfaces projected onto the *R̅*–SJL plane. The x-axis represents *R̅*, the y-axis represents SJL. Filled contours indicate the predicted GPA levels, while contour lines highlight gradients of change. Finches show the highest predicted GPA at moderate levels of both *R̅* and SJL, forming a smooth, elliptical peak across the surface. Among Larks, GPA varies primarily with *R̅*, displaying a clear quadratic pattern, while SJL has little to no influence on performance. Owls exhibit a trend-level interaction between *R̅* and SJL. When SJL is minimal, Owls with low circadian distinctness achieve the highest predicted GPA of any group, challenging the common assumption that evening types inherently perform worse academically. In contrast, when SJL is extreme, owls with higher *R̅* show slightly improved grades relative to those with low *R̅*, indicating that the costs or benefits of circadian distinctness depend strongly on individuals’ alignment, or misalignment, with external schedules. Abbreviations: R - *R̅*, distinctness of the circadian rhythm, SJL - social jetlag, GPA - grade point average.

The variance explained by the models was small but consistent across methods, ranging from 1-4% depending on chronotype, with owls showing the strongest association (*R̅*^2^ ≈ 3-4%). Details are provided in Supplementary table S17.

## Discussion

This study shows that the distinctness of an individual’s circadian rhythm – captured through the concentration (*R̅*) of their behavioral activity across the 24-hour cycle – is meaningfully related to academic performance. Using a large, real-world dataset of over 3.4 million LMS login events from 13,894 students, we demonstrate that circadian distinctness and social jetlag jointly shape educational outcomes and that these relationships differ systematically across chronotypes. To our knowledge, this is the first study to demonstrate that the distinctness of the circadian rhythm plays an important role in academic performance. Together, our findings highlight the multidimensional nature of human circadian functioning and offer an objective behavioral approach to quantifying a construct that has previously relied almost entirely on self-report questionnaires.

Across the student population, moderate levels of distinctness were associated with the highest grades, suggesting that both highly rigid and highly dispersed daily activity patterns may hinder academic performance. This observation remained consistent across semesters, seasons, and sex (Fig. 1). This consistency suggests that *R̅* is a stable behavioral trait, relatively unaffected by typical demographic or environmental covariates. Yet this overall pattern masked substantial chronotype-specific differences (Fig. 2). Morning types performed better with higher distinctness, consistent with the idea that strong alignment between internal timing and early institutional schedules confers an academic advantage. Intermediate types showed the highest performance at moderate distinctness, while evening types displayed the opposite pattern – high distinctness was associated with lower grades. These results challenge the widespread assumption that all evening types are uniformly disadvantaged. Our analyses reveal that only evening types with rigid daily rhythms perform poorly, those with more flexible (low-distinctness) rhythms achieve grades comparable to, or even exceeding, morning types.

Distinctness was also systematically related to social jetlag. Students with more concentrated activity patterns generally showed larger shifts between class days and non-class days. This is consistent with theoretical assumptions behind the concept of high distinctness - individuals with very sharply defined rhythm tend to adjust their diurnal functioning to externally imposed schedules less effectively, and in free days, they often return to their natural, endogenous biological rhythm. This pattern was stable across semesters, seasons, and sexes (Fig. 3). However, chronotypes again differed: intermediate types showed a monotonic increase in social jetlag as distinctness rose, while morning and evening types exhibited an U-shaped relationship (Fig. 4). This suggests that both very rigid and very diffuse activity patterns in Owls may promote misalignment with institutional schedules, although likely through different behavioral mechanisms. We should also note that owls, as a group, exhibit substantially higher levels of social jetlag^25^. If larks and finches experienced comparable SJL levels their patterns might similarly converge.

Finally, when distinctness and social jetlag were considered jointly, the students with the highest predicted grades were evening types who showed low distinctness and minimal social jetlag. Conversely, larks appeared less vulnerable to social jetlag than Finches and Owls, as rhythm distinctness contributed more strongly to their GPA. Finches, however, performed best under conditions of minimal social jetlag and moderate distinctness (Fig. 5 and Fig. 6). These findings clarify that the academic disadvantage often attributed to eveningness does not stem from evening preference itself, but from the overall interplay of chronotype, distinctness, and schedules that are incompatible with the students’ biological timing.

Our interpretation is consistent with prior psychological and neurobiological findings. Questionnaire-based studies have shown that individuals with elevated circadian distinctness display reduced conscientiousness^28,29^ (which negatively affects academic performance^31,32^), greater neuroticism, avoidance behavior, sensitivity to punishment and negative emotionality^14,29,30,33–35,40^. Neuroimaging findings similarly indicate that circadian distinctness may play an important role in shaping brain structure^36^ and task-related activity ^41^, particularly in regions responsible for cognitive processes, like attention, semantic processing, working memory, and executive functions^42–46^. Together with the present results, these findings suggest that circadian distinctness is not merely a subjective feeling but reflects a broader scope of cognitive and neurobiological characteristics that may impact real-world academic performance.

It is important to note that in circular statistics, the mean resultant length (*R̅*) is traditionally interpreted as a measure of concentration – how tightly clustered observations are around a preferred phase^22^. Precisely the aspects that *R̅* captures mathematically. For this reason, we proposed that *R̅* is a promising objective behavioral marker of circadian distinctness, particularly in large datasets such as LMS logins. However, we currently do not possess a dataset in which subjective questionnaire-based distinctness and objective *R̅*-based distinctness can be directly compared. Thus, while our interpretation is theoretically grounded, we cannot yet claim that mean resultant length (*R̅*) constitutes the definition of circadian distinctness. Further studies combining behavioral time-series with questionnaires will be essential for formally establishing equivalence. Within the constraints of the present dataset, *R̅* represents the best available and conceptually well-justified measure of distinctness.

Our dataset offers unprecedented ecological validity. However it presents some challenges. This dataset captures real human behavior in authentic contexts, not biased by laboratory conditions. While this increases generalizability, it limits the possibility of extracting information about other daily obligations or constraints of students that may influence login timing (personal constraints, lifestyle, workload, etc.). Thus, some activity patterns may reflect availability rather than internal circadian preference. Furthermore, LMS logins are inherently irregular: students differ widely in frequency and timing of activity. This irregularity limits the use of traditional chronobiological tools such as cosinor or periodograms, which require more structured time series (many of their assumptions are violated in these real-life conditions). Therefore, we paid special attention to implement strict filtering and correct preprocessing to ensure that the analyzed students exhibited sufficiently informative activity patterns for circadian inference. Moreover, the exceptionally large sample size substantially reduced the impact of these limitations.

Although the variance in academic performance explained by distinctness, social jetlag and chronotype was modest (approximately 4%), this magnitude is typical for educational datasets ^26,47–49^ and is far from trivial, given the diverse and uncontrolled influences on student performance. These effects, while small at the individual level, may accumulate meaningfully across populations, particularly in institutions that rely heavily on early morning schedules.

Overall, our results have important societal implications and demonstrate that improving educational equity may require moving beyond traditional chronotype-based categorizations and considering the flexibility (or rigidity) of students’ daily rhythms to avoid one-size-fits-all schedules.

## Conclusions

In this study, we explore a previously overlooked dimension of human circadian rhythmicity: circadian distinctness (subjective amplitude). Distinctness represents how strongly an individual’s daily rhythm fluctuates across the day. Using Circular statistics, we propose the first objective behavioral measure of distinctness and show that it is a promising predictor of academic performance. Students with moderate rhythmicity performed best overall, but this pattern varied strikingly by chronotype: larks benefited from stronger rhythms, finches from moderate rhythms, and owls from weaker, more flexible rhythms. Circadian distinctness was also closely linked to social jetlag, which increased with more rigid rhythmicity across chronotypes. When both factors were considered together, low social jetlag and moderate distinctness were associated with the highest academic performance. Our analyses further reveal that not all owls are the same: only evening types with rigid, highly distinct daily rhythms show poor academic outcomes. Evening types with more flexible, low-distinctness rhythms achieve grades comparable to, or even exceeding, those of morning types. Together, these results establish circadian distinctness as a meaningful and previously neglected dimension of human rhythmicity, highlighting the importance of circadian flexibility for academic success, and underscoring that uniform early schedules may disproportionately disadvantage students whose biological rhythms run later.

## Materials and methods

### Data collection

Under the Northeastern Illinois University institutional review board (IRB) protocol #16-073 MO1, data from 13,894 students were collected, de-identified, and processed as described previously^25^ In this work we focus solely on “non-class days”, as in our previous work we showed that login times from “class days” mimic the actual class schedule, making it impossible to distinguish between each student’s endogenous (biological internal rhythmicity) and exogenous (driven by external cues) rhythm. Individual semester GPAs were calculated by converting letter grades into their numerical equivalent grade points (A=4.0), which were then averaged for each student each semester. All analyses and visualizations were performed using Python libraries NumPy 1.26.4, Pandas 2.2.3, Statsmodels 0.14.4, Scikit-learn 1.1.3, and Matplotlib 3.9.2

### Circadian characteristics

In this study, to obtain a formal measure of circadian rhythm distinctness, we employed the mean resultant length *R̅*, calculated from students’ login times on non-class days. The concentration is computed directly from login times represented as unit vectors on the 24-hour clock. This metric quantifies how concentrated or dispersed activity is throughout the day and is calculated as the magnitude of the resultant vector (*r*), i.e. the vector sum of all clock-time vectors ^20,50,51^. The resultant length (*R* = |*r*|), divided by the number of contributing vectors (*n,* here: logins) yielding the mean resultant length (*R̅*). The value *R̅* ∈ [0,1] and it approaches 1 if and only if the sample is tightly clustered, while it approaches 0, if the sample is widely dispersed or antipodally symmetric.

In the context of circadian rhythmicity, *R̅* value of 0 indicates that logins are distributed broadly across the full 24-hour day, whereas *R̅* value of 1 would indicate that all logins occurred at precisely the same time each day. We interpret higher *R̅* as greater regularity or stability of the individual’s circadian rhythm: if a student logs into the system at approximately the same time each day, the “window” of activity is narrow, and the distinctness of the rhythm is high. Conversely, if a student logs in at different times each day with little regularity, the activity window is broad, and the distinctness of circadian activity is low.Schematic representation of this concept and exact equation for *R̅* calculations are presented in Supplementary materials S1, S2.

In our previous work, clock time values from the LMS were used to derive circular time-as-radians (0 to 2π radians) measures for each login event, with a circular period of 24 hours. Median radial login phase was employed to define individual chronotype and social jetlag. All individuals within each semester were classified as a lark, finch, or owl. Larks were defined as those with median non-class day phases from more than one standard deviation below the group median to pi radians from that median. Owls were defined as those with median non-class phases from more than one standard deviation above the group median to pi radians later than that median. Individuals within one standard deviation of the group median were designated as finches. This approach led to the identification of 13372 Finches, 1419 Owls, and 2417 Larks. To confirm the correctness of previously defined chronotype, we additionally computed mean direction, µ. The formula for calculating the mean direction is provided in Supplementary materials S1, S2. Social jet-lag was defined by subtracting the non-class day median phase from the class day median phase for each individual, yielding a number of hours of median phase-shift from non-class days to class day as described previously ^25^. Because students showed both delayed and advanced shifts between non-class and class days, we observed positive and negative social jetlag across the sample.

### Data preprocessing

We preprocessed and filtered our data prior to our statistical analyses. Because students were free to decide if and when to log into the system, the dataset exhibited substantial heterogeneity in sampling rate and uneven temporal spacing, violating assumptions required for standard circadian methods such as cosinor or periodogram analyses. To address these characteristics, we followed the workflow described by Karoly and colleagues^52^ who analyzed similarly irregular behavioral-event data.

We first assessed whether students’ login times were uniformly distributed across the 24-hour cycle. Three complementary tests were applied: (I) Rayleigh test, which evaluates uniformity under the assumption of a unimodal von Mises distribution (the circular analogue of the Gaussian distribution), (II) Watson U^2^ test which assesses goodness-of-fit to von Mises assumptions, and (III) the Hodges-Ajne test, which is distribution-free and can identify the unimodal, bimodal, and multimodal distributions. To account for multiple comparisons across individual-level tests, *p*-values from the Rayleigh, Watson U^2^, and Hodges-Ajne tests were corrected for multiple tests using the Benjamini-Hochberg false discovery rate (FDR) procedure (q < 0.05) within each academic term.

Because the mean resultant length (*R̅*) is known to be biased upward for small sample sizes and its variance increases when the true concentration is low^53^, we used a Monte-Carlo simulation to derive distribution-free significance thresholds for *R̅* under circular uniformity (to assess how many logins are sufficient to obtain a significant *R̅* value). For each possible number of logins (*n*) observed across students from our dataset, we generated 10,000 random samples of *n* angles (login times) drawn from a uniform [0, 2π) distribution, and computed their corresponding *R̅* values. The 95^th^ and 99^th^ percentiles of these simulated *R̅* distributions served as critical values, against which each student’s observed evaluated. *R̅* was evaluated.

To ensure stable inference and to avoid artificial inflation of *R̅* due to small *n*, we identified the minimum number of logins per student required to have robust *R̅* value. To ensure stable inference from the simulation-based *R̅* thresholds, we identified the smallest logins number *n* at which the null threshold curves (95th/99th percentiles under uniformity) became effectively flat (Figure in Supplementary Materials S3). We computed gradients d*R̅*/d*n* and defined the stabilization point as the smallest *n* with d*R̅*/d*n* < 0.005. In other words, we detected the point (*n*_stable_) where each curve flattens out – where the change in threshold per additional login falls below 0.005. Further analyses of *R̅* were restricted to students with *n* ≥ *n*_stable_. The 99^th^ percentiles curve flattened at ∼26 logins, indicating that above this point the number of logins no longer artificially inflates *R̅*. Subsequent analyses were restricted to students with n ≥ 26 logins per term. Full details of the simulation procedure and threshold derivation are provided in Supplementary materials S3.

### Statistical analysis

#### Group differences in ***R̅***

Normality of the *R̅* values were assessed using Shapiro-Wilk (for sample size <5000) and D’Agostino-Pearson tests (for sample size >5000). Sample sizes for each semester and chronotype are reported in Supplementary Table S6. To test for differences in *R̅* between semesters, sex and chronotype groups, we used nonparametric tests. Overall group effects were assessed using Kruskal-Wallis tests, followed by pairwise Mann-Whitney U comparisons with FDR multiple comparison corrections. Effect sizes (ε^2^ for Kruskal-Wallis and |r| Mann-Whitney U) were computed to quantify the magnitude of group differences.

#### Relationship between ***R̅***, SJL and GPA

We modeled GPA as a function of mean-centered *R̅* (*R̅*_cent_) using Generalized Estimating Equations (GEE) to account for repeated semesters within students. The model included linear and quadratic terms to capture potential nonlinearity. We first fit a base model including only *R̅*_cent_ , *R̅*^2^ . Next, we checked whether the relationship between *R̅* and GPA is moderated by season, semester and sex by adding interaction terms in follow-up GEE analyses. To check if shape of the *R̅*-GPA relationship differed between chronotypes, we fitted GEE models separately within each chronotype group. We visualized all these relationships using locally weighted regression (Locally Estimated Scatterplot Smoothing, LOESS) fitted separately by term, by season, by sex and by chronotype. Curves were smoothed with a span of 0.6 and accompanied by pointwise 95% bootstrap confidence bands (1000 iterations). Details of the models are provided in the Supplementary materials. Our previous paper demonstrated that SJL is a significant predictor of GPA, thus in present study we additionally examined the relationship between *R̅* and SJL as well as joint association between *R̅* and SJL with GPA. GPA was modeled as a function of the linear terms for *R̅* and SJL, their quadratic components, and their higher-order interaction. To visualize the joint association between *R̅* and SJL with GPA, we generated 3D prediction surfaces using generalized estimating equation (GEE) models, in which the x-axis represents *R̅*, the y-axis represents SJL (in hours), and the z-axis displays the predicted GPA. To provide a complementary 2D interpretation of the same GEE-predicted surfaces, we generated contour maps, which display predicted GPA values across the *R̅*–SJL plane, while overlaid contour lines mark isolines of equally predicted GPA. Finally, to assess whether the joint association of *R̅* and SJL with GPA differed by chronotype, we fitted the same GEE model separately in each chronotype group and visualized these relationships using both 3D and 2D contour plots. To determine whether a linear or quadratic specification provided a better fit within each chronotype group, we compared GEE models using the Quasi-Information Criterion (QIC), following Pan^39^. Effect size was assessed using Zhang’s pseudo-R² and correlation-based pseudo-R² adapted for marginal models.

## Acknowledgments

The authors would like to thank the Northeastern Illinois University Office of Institutional Research and Assessment and the Center for Teaching and Learning for their assistance in organizing the original data set. We would also like to thank Dr. Benjamin L. Smarr, who co-authored the first publication using these data and whose chronotype classification was used in the current study, for his continued collegial support. We are also grateful to all the students whose data were analyzed in this study.

## Code and data availability statement

Custom Python scripts as well as summarized LMS data used in this work are available freely in the GitHub repository: https://github.com/PatrycjaScislewska/LMS_Distinctness_of_the_circadian_rhythm

## Authors contributions

All authors conceived the project. All authors interpreted the results and revised the manuscript. P.S.: Methodology, Formal analysis, Code development, Writing - Original Draft, Visualization; P.B.: Funding acquisition, Writing - review & editing, Supervision; I.S.: Writing - review & editing, Supervision; R.S.: Writing - review & editing, Supervision; A.E.S. Conceptualization, Study design, Data collection, Methodology, Formal analysis, Code development, Funding acquisition, Writing - review & editing, Supervision

## Supplementary Materials

### S1. Equation for calculation of mean direction, **µ**

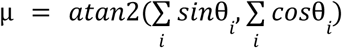

where

θ – angle *i* in radians, representing the time of login *i*

### S2. Equation for calculation of mean resultant length, ***R̅***

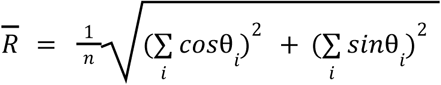

### where

θ – angle *i* in radians, representing the time of login *i*

n – number of logins

**Visual, schematic representation of Mean Direction (marked with orange arrow) and Mean Resultant Length.**

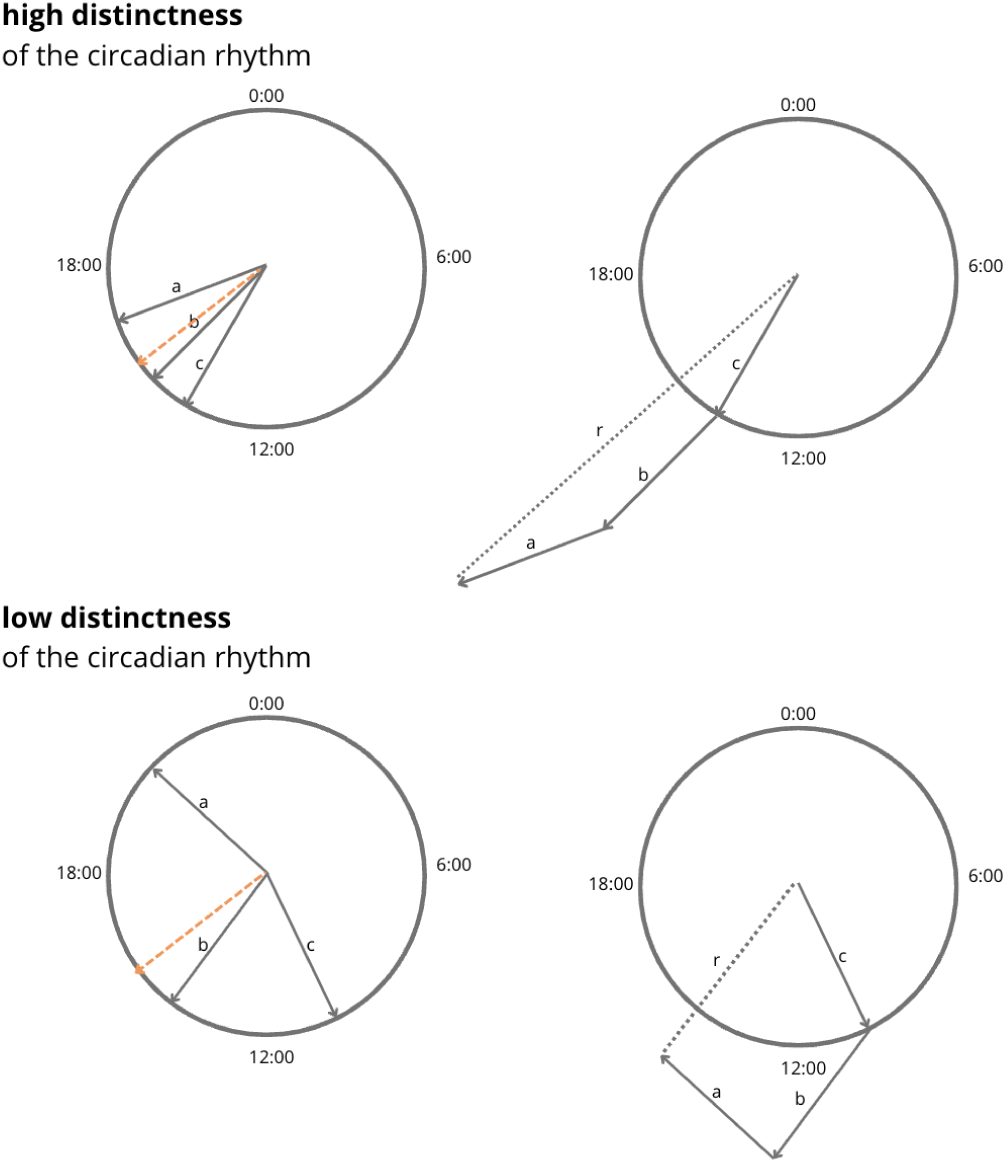

To compute the mean resultant length we place all of the vectors (login times in radians) head to toe. The length of the resulting new vector is the resultant length. The mean resultant length, *R̅*, is the length of this vector divided by the number of vectors from which it was created. *R̅* differs between 0 and 1. In this example, right activity is more concentrated, resulting in smaller *R̅*, while left activity is more spread our, resulting in greater *R̅*.

**The correspondence between concept of the distinctness of the circadian rhythm and Mean Resultant Lenght**

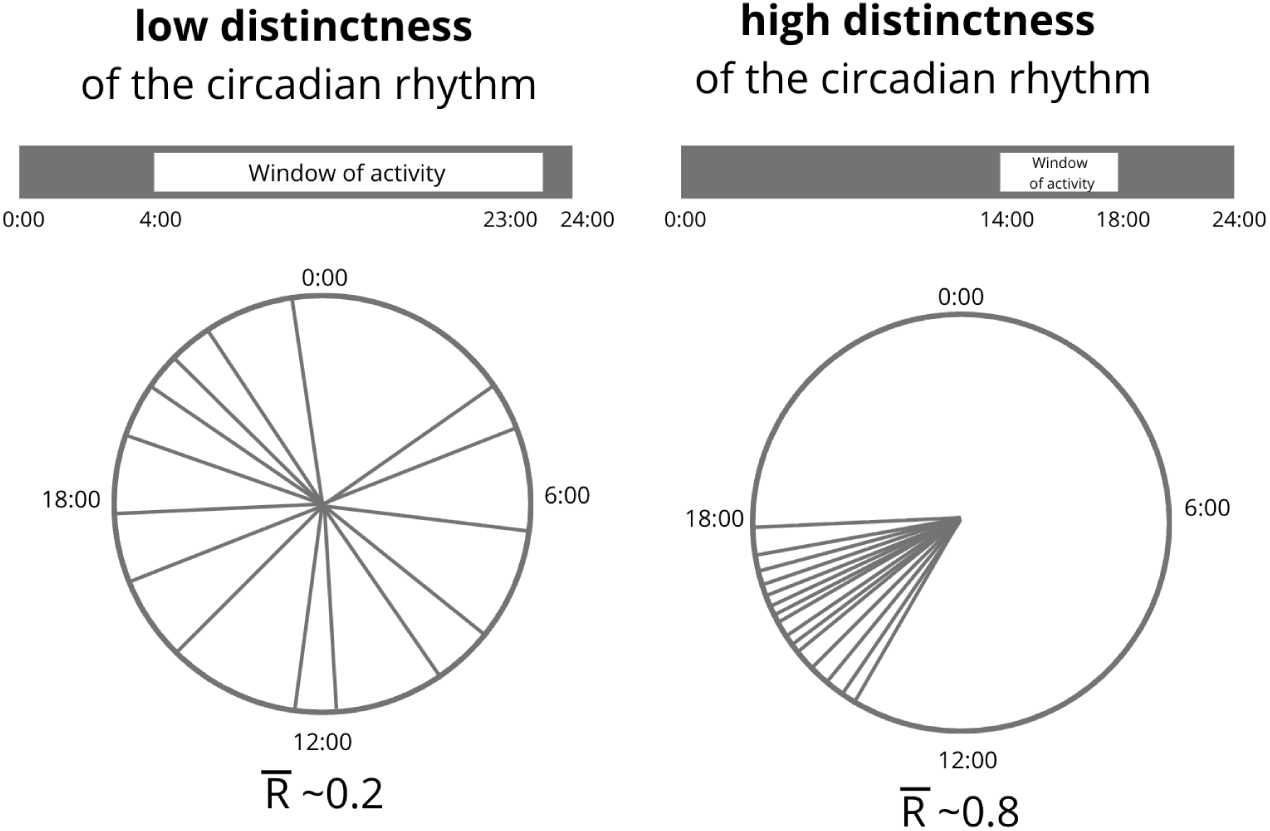

### S3. Students’ logins data distributions

Pattern labeling was based on circular uniformity tests as follows: participants with fewer than two valid observations were excluded. For the remaining participants, if the Hodges-Ajne test indicated a significant deviation from uniformity (*p* < 0.05), the pattern was classified as non-uniform, within these, a significant Rayleigh test (*p* < 0.05) defined the pattern as unimodal, whereas a non-significant Rayleigh test defined it as multimodal. If the Hodges-Ajne test was not significant, the pattern was classified as uniform. To account for multiple comparisons, the Benjamini-Hochberg false discovery rate (FDR) procedure (q < 0.05) within each academic term was applied.

Circular statistical tests consistently revealed significant circadian rhythmicity in students’ activity across all semesters (Rayleigh *p* < .05 for 78.3% of cases, Watson *p* < .05 for 80.8%). Most students displayed a unimodal pattern of activity (69.3%), while only 2.9% showed multimodal distributions and 27.8% uniform patterns. The proportion of students exhibiting significant rhythmicity remained stable across semesters, with 65.5-75.7% showing non-uniform daily patterns according to Rayleigh, Hodges-Ajne, and Watson tests.

| Term | Students | Valid Students (with > 2 logins) | Insufficient Students (with $\leq 2$ logins) | Rayleigh $p < .05$ | Hodges-Ajne $p < 0.05$ | Watson $p < 0.05$ | Unimodal | Multimodal | Uniform |
| --- | --- | --- | --- | --- | --- | --- | --- | --- | --- |
| Fall 2014 | 8,558 | 8,192 | 366 | 6,144 (75.0%) | 5,652 (69.0%) | 6,343 (77.4%) | 5,127 (62.6%) | 235 (2.9%) | 2,830 (34.5%) |
| **Spring 2015** | 7,925 | 7,711 | 214 | 6,102<br>(79.1%) | 5,880<br>(76.3%) | 6,314<br>(81.9%) | 5,500<br>(71.3%) | 204 (2.6%) | 2,007<br>(26.1%) |
| **Fall 2015** | 8,795 | 8,574 | 221 | 6,790<br>(79.2%) | 6,526<br>(76.1%) | 6,975<br>(81.4%) | 6,071<br>(70.8%) | 249 (2.9%) | 2,254<br>(26.3%) |
| **Spring 2016** | 8,051 | 7,841 | 210 | 6,276<br>(80.0%) | 6,102<br>(77.8%) | 6,489<br>(82.8%) | 5,691<br>(72.6%) | 243 (3.1%) | 1,907<br>(24.3%) |
| **All terms** | 33,329 | 32,318 | 1,011 | 25,312<br>(78.3%) | 24,160<br>(74.8%) | 26,121<br>(80.8%) | 22,389<br>(69.3%) | 931 (2.9%) | 8,998<br>(27.8%) |

### Monte-Carlo simulation

For each student, the spread of the logins was quantified as the mean resultant length *R̅*. Because small sample sizes can yield spuriously high *R̅* values even under uniformity and the expected value of *R̅* depends on the number of observations a student has, we estimated the null distribution of *R̅* (the distribution expected if logins were completely random over 24 hours) using Monte-Carlo simulation.

Using a fixed pseudorandom seed for reproducibility, we generated *N*_sims_=10,000 random samples of size *n* drawn from Uniform [0, 2π) for a grid of *n* values covering both the empirical range and additional reference points (5, 10, 20, 30, 40, 50, 75, 100, 150, 200). For each *n*, the mean resultant vector length, defined by the formula provided in S2, was calculated for every simulation. In other words, for each possible number of logins *n*, we generated 10,000 samples of *n* random times drawn from a uniform circular distribution and calculated the corresponding *R̅* values.

The 95^th^ and 99^th^ percentiles of the resulting null distributions defined critical thresholds, which serve as significance thresholds for detecting non-uniformity. An observed *R̅* above these thresholds indicates that the student’s login behavior is more clustered in time than expected by chance.

Each student’s observed *R̅* was compared to interpolated values of R_95_(*n*) and R_99_(*n*) corresponding to their actual number of events. Values exceeding these thresholds were marked as significant at the 5% or 1% level, respectively. Threshold curves and per-term scatterplots (*R̅* vs *n*, log-scaled x-axis) were visualized to illustrate how *R̅* relates to sample size.

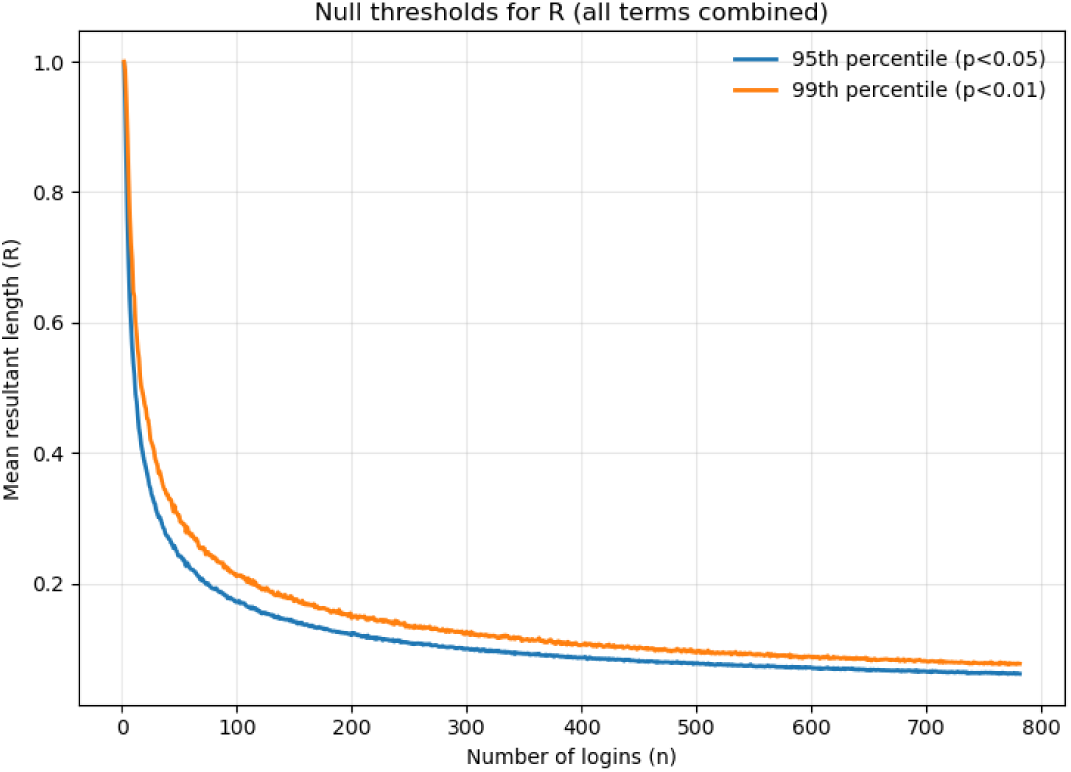

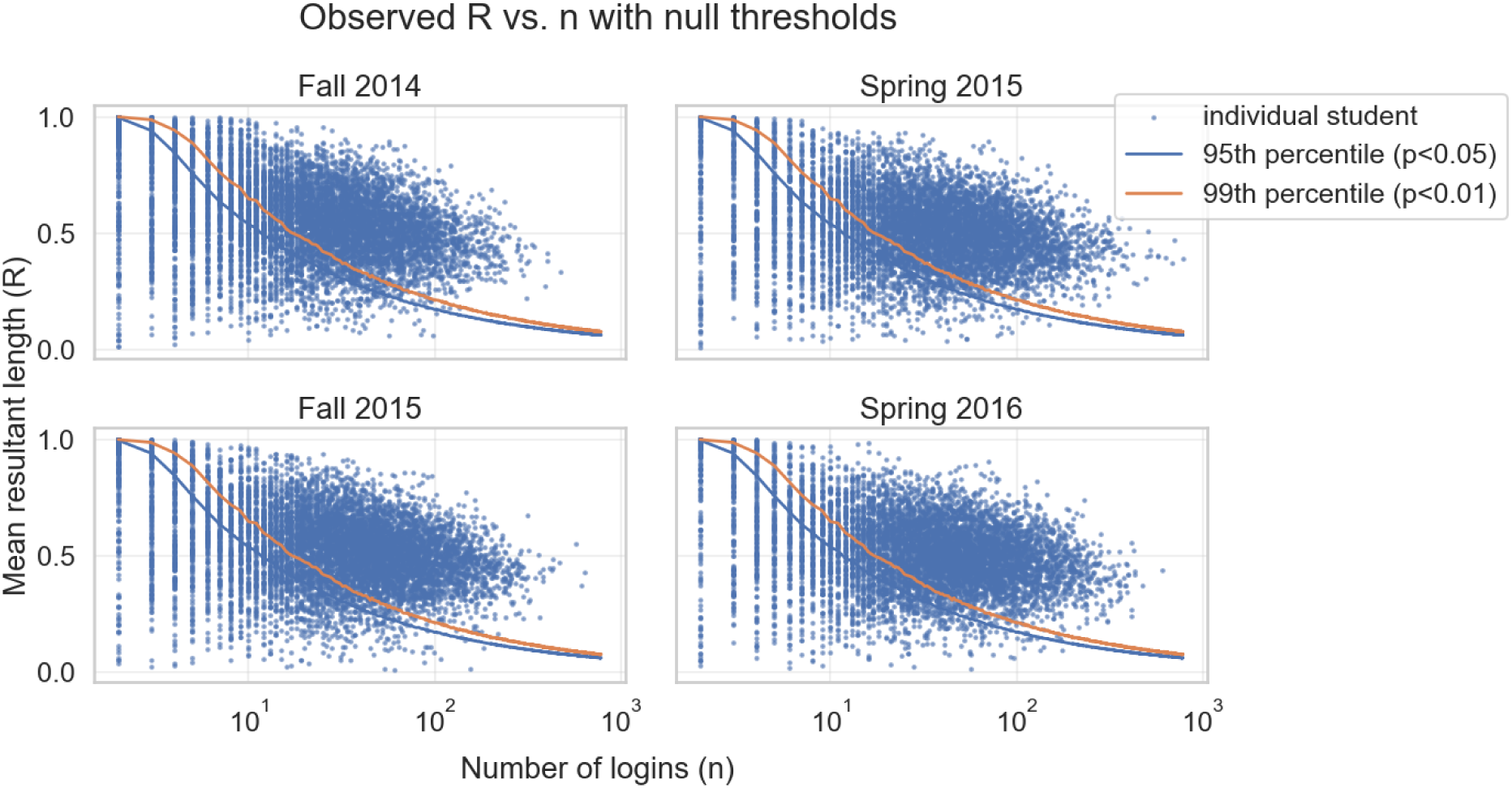

The simulation-derived null thresholds R_95_(*n*) and R_99_(*n*) change across different numbers of logins – decrease with sample size and approach an asymptote. To define a stabilization point for each curve, we computed numerical gradients dR/dn. For each curve, the stabilization point *n*_stable_ was the smallest *n* such that the absolute gradient fell below 0.005.

We adopted a conservative filter by requiring each student has number of logins *n* ≥ *n*_stable_ . Based on the 99% curve, we found that *n*_stable_ is equal to 26. Therefore, to ensure that R was estimated reliably and not biased by small sample sizes, we restricted all analyses to students with at least 26 login events. For transparency, we plotted R_95_(*n*) and R_99_(*n*) against *n* with vertical lines at the corresponding stabilization points and reported per-term retention rates after applying the minimum-*n* filter.

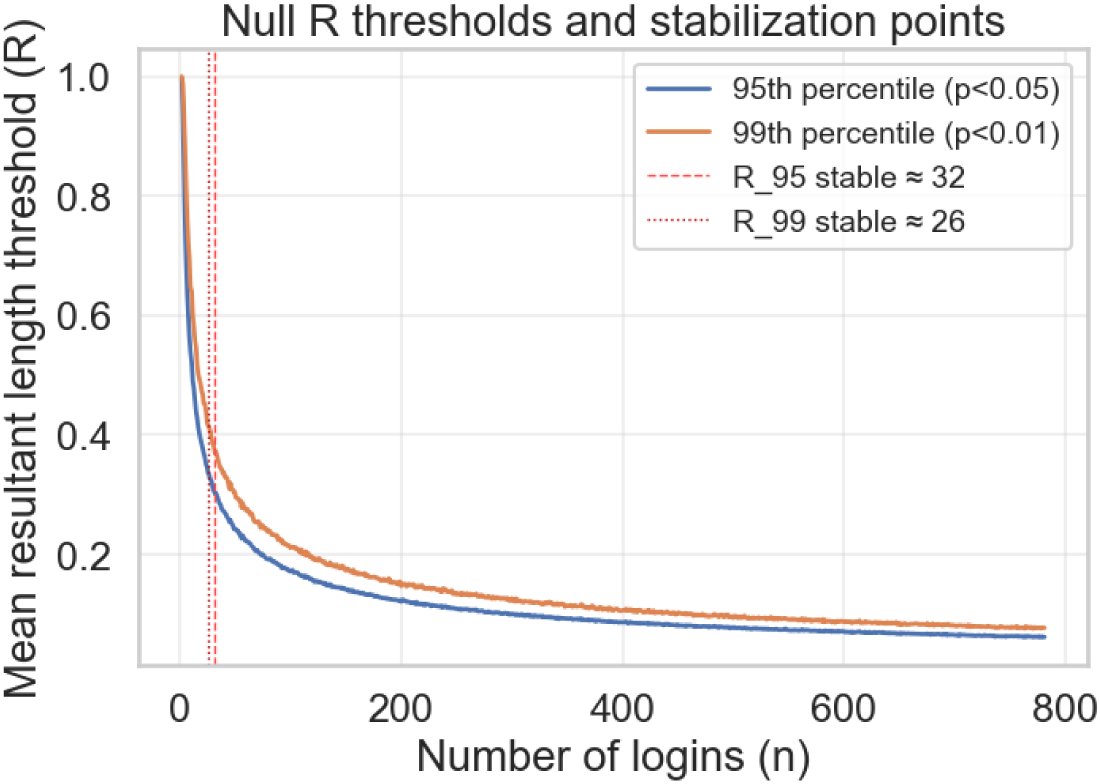

Fall 2014: kept 3572/8558 (41.7%) with *n* ≥ 26

Spring 2015: kept 4379/7925 (55.3%) with *n* ≥ 26

Fall 2015: kept 4737/8795 (53.9%) with *n* ≥ 26

Spring 2016: kept 4688/8051 (58.2%) with *n* ≥ 26

This simulation-based threshold guarantees no artificial inflation of R by small sample size.

Importantly, we removed students who had less than 26 logins per term, but we did not remove the students who are below the simulation-based null threshold curves, as we do not require students to be rhythmic. Students with n ≥ 26 may still have low or non-significant R scores, which simply reflects genuinely irregular or uniform login behavior rather than measurement noise.

To have the complete characterization of our dataset after filtering, we checked the distributions of logins (uniform / non-uniform distributions) of students remaining after filtering. Among the 17,376 students who met criterion of at least 26 logins, the large majority showed significant non-uniformity (Rayleigh *p* < .05: 96.2%; R > R₉₅: 96.2%; R > R₉₉: 91.6%), indicating that most of students with primarily uniformly distributed logins were in fact the effect of too sparse login density and in our final dataset most students have non-uniform distribution of logins. The nearly perfect overlap between the Rayleigh test and simulation-based thresholds confirms the robustness of our filtering simulation based method and confirms the circadian characteristics of activity of students. This is consistent with results presented in our previous paper ^25^.

| Metric | Count | Percentage |
| --- | --- | --- |
| Total students ($n \geq 26$) | 17,376 | 100 % |
| Rayleigh test $p < 0.05$ | 16,708 | 96.2% |
| $R > 95\%$ simulation-based threshold | 16,711 | 96.2% |
| $R > 99\%$ simulation-based threshold | 15,919 | 91.6% |
| Overlap: Rayleigh $p < .05$ AND $R > 95\%$ | 16,704 | 96.1% |
| Overlap: Rayleigh $p < .05$ AND $R > 99\%$ | 15,919 | 91.6% |

### S4. Descriptive statistics – dataset after filtering

|  | n_events | R_bar | mu_rad | mu_hours | TERM_GPA |
| --- | --- | --- | --- | --- | --- |
| count | 17376. | 17376 | 17376 | 17376 | 17376 |
| mean | 69.865677 | 0.493451 | 4.388697 | 16.763589 | 3.260915 |
| std | 52.432248 | 0.133647 | 0.641974 | 2.452161 | 0.796442 |
| **min** | 26.000000 | 0.008706 | 0.032320 | 0.123453 | 0.000000 |
| **25%** | 37.000000 | 0.405162 | 3.973467 | 15.177527 | 3.000000 |
| **50%** | 53.000000 | 0.494851 | 4.419266 | 16.880351 | 3.500000 |
| **75%** | 83.000000 | 0.582247 | 4.845210 | 18.507339 | 4.000000 |
| **max** | 781.000000 | 0.941884 | 6.245514 | 23.856105 | 4.000000 |

### S5. Rose plots - distribution of mean hours of activity within groups of chronotypes

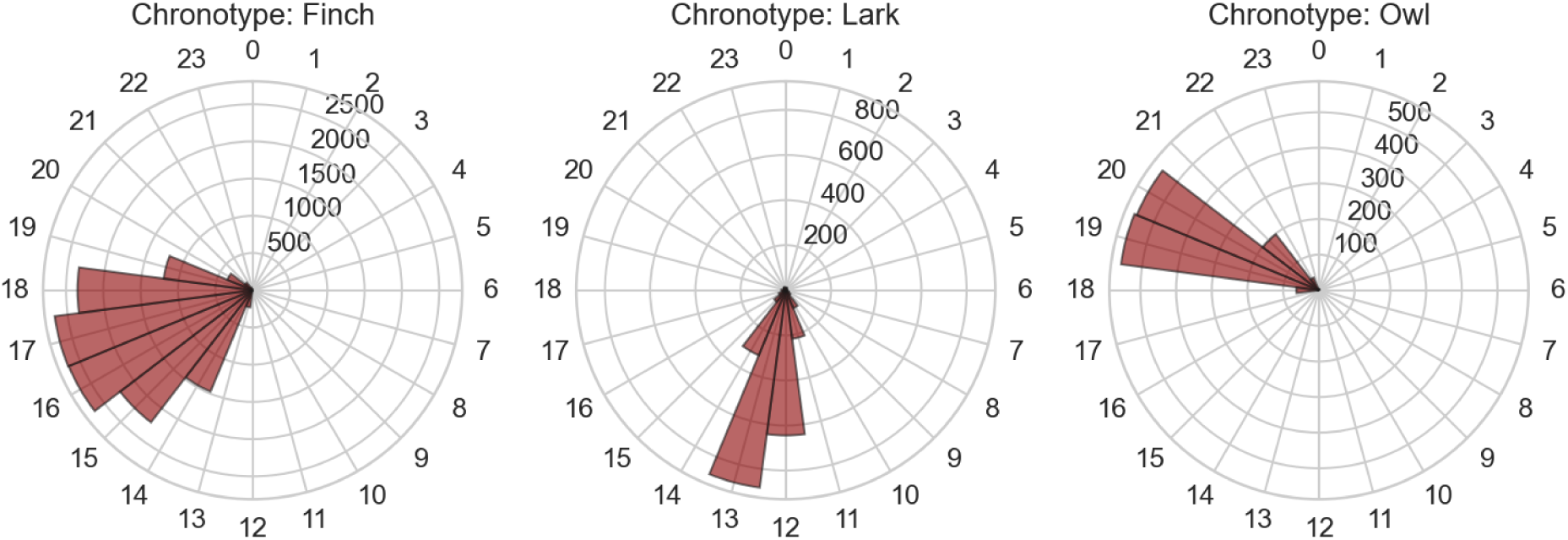

| Category | Mean ($\mu$, rad) | Mean ($^{\circ}$) | Mean (hours) | N (students) |
| --- | --- | --- | --- | --- |
| **Overall** | 4.41 | 252.5$^{\circ}$ | 16.84 h | 17376 |
| **Fall 2014** | 4.46 | 255.7$^{\circ}$ | 17.05 h | 3572 |
| **Spring 2015** | 4.39 | 251.5$^{\circ}$ | 16.76 h | 4379 |
| **Fall 2015** | 4.44 | 254.5$^{\circ}$ | 16.96 h | 4737 |
| **Spring 2016** | 4.35 | 249.2$^{\circ}$ | 16.62 h | 4688 |
| **Chronotype = Finch** | 4.46 | 255.7$^{\circ}$ | 17.05 h | 13 372 |
| **Chronotype = Lark** | 3.45 | 197.7$^{\circ}$ | 13.18 h | 2 417 |
| **Chronotype = Owl** | 5.30 | 303.4$^{\circ}$ | 20.23 h | 1 419 |

### S6. Test of normality of ***R̅*** values

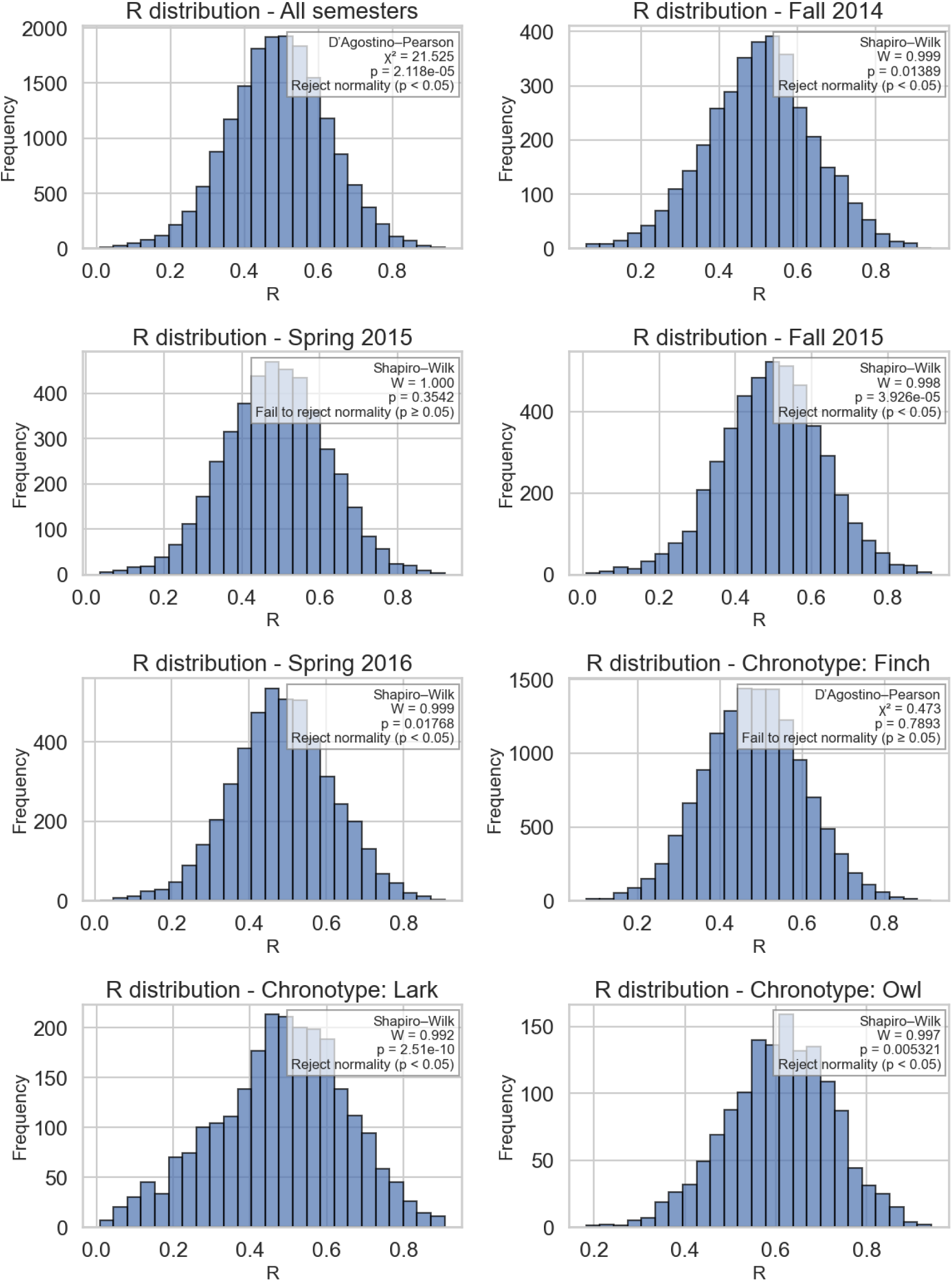

### S7. Group differences in R - pairwise comparison between terms and seasons

Group differences in *R̅* between semesters, sex and chronotypes, were assessed using nonparametric Kruskal-Wallis test and followed by pairwise Mann-Whitney U tests with FDR multiple comparisons correction. Seasonal differences (both falls and both springs combined) were statistically significant (p < 0.05), whereas comparisons within seasons (fall 2014 vs fall 2015, spring 2015 vs spring 2016) were not. The effect of season was very small (ε^2^ = 0.003), and sex differences were even smaller (ε^2^ = 0.001). With a dataset of over 17,000 students, even tiny differences can yield statistically significant p-values because the standard error decreases with √n. Thus, these negligible effect sizes suggest that the significant results for season and sex are driven by the large sample size rather than by meaningful differences.

In contrast, chronotype differences were significant and showed a larger effect (ε² = 0.065), indicating that chronotype explains about 6.5% of the variability in circadian distinctness. Owls showed higher *R̅* values than Finches and Larks, meaning that Owls are substantially more consistent in their login timing than Finches or Larks.

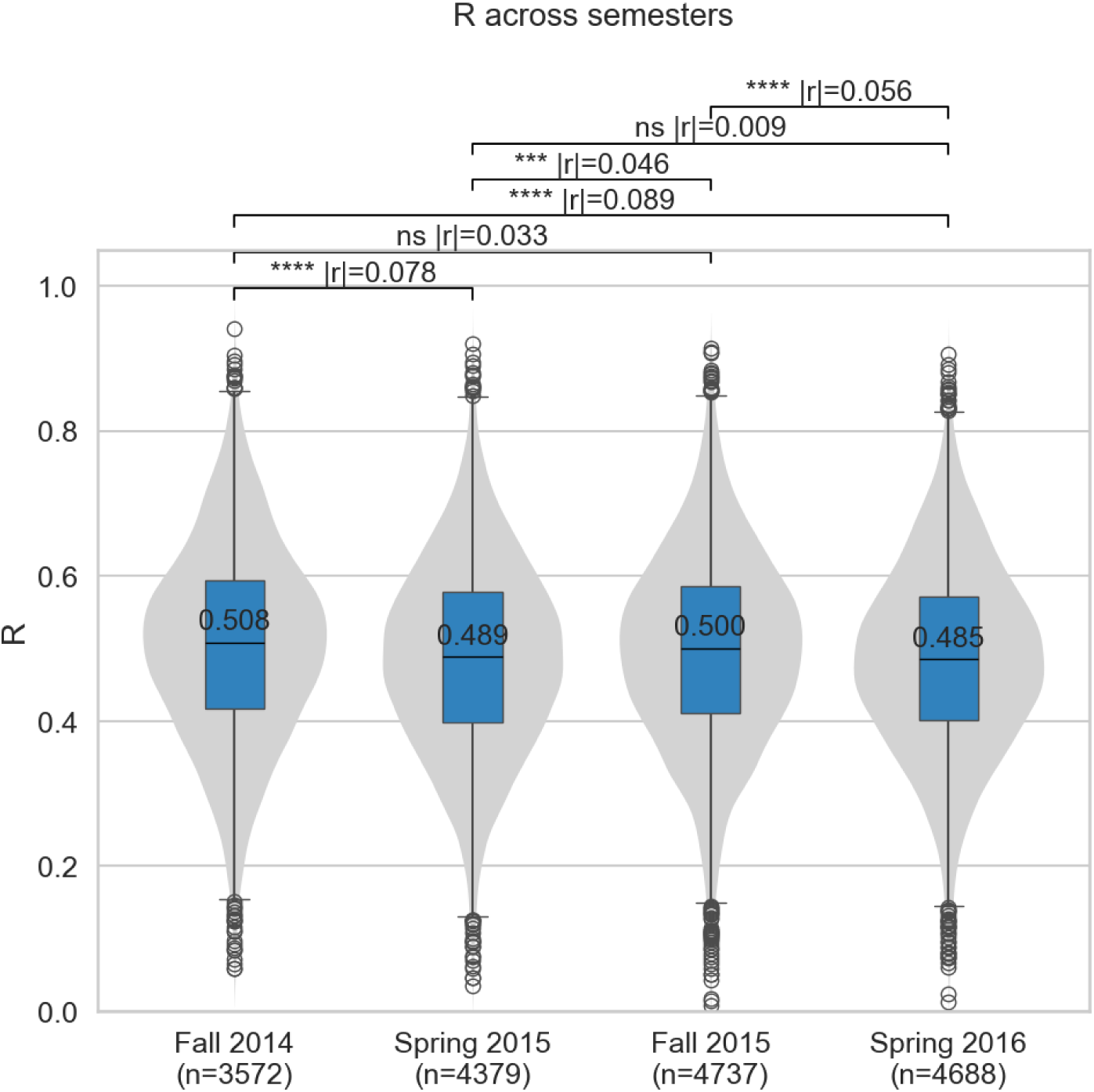

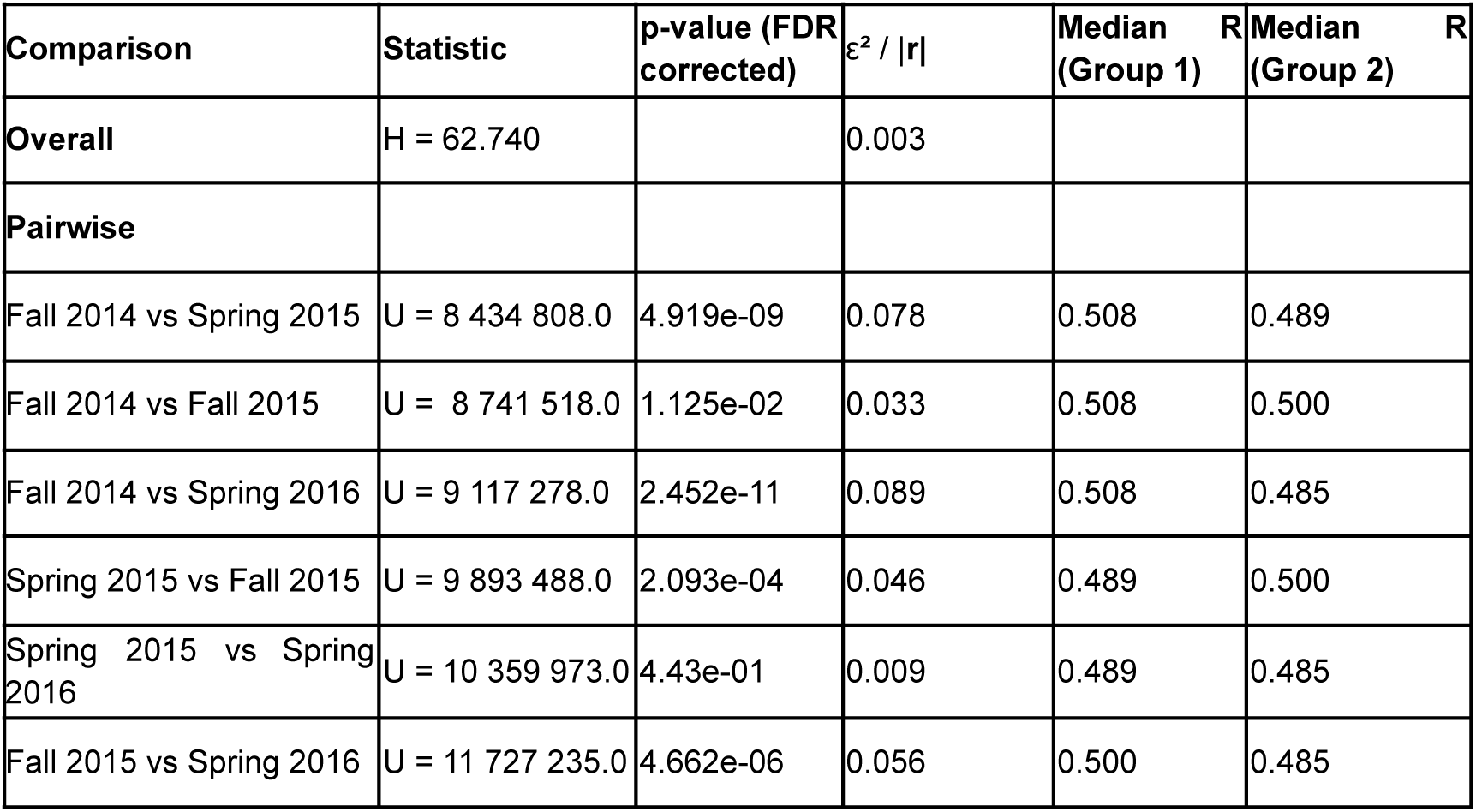

### S8. Group differences in R - pairwise comparison between sex

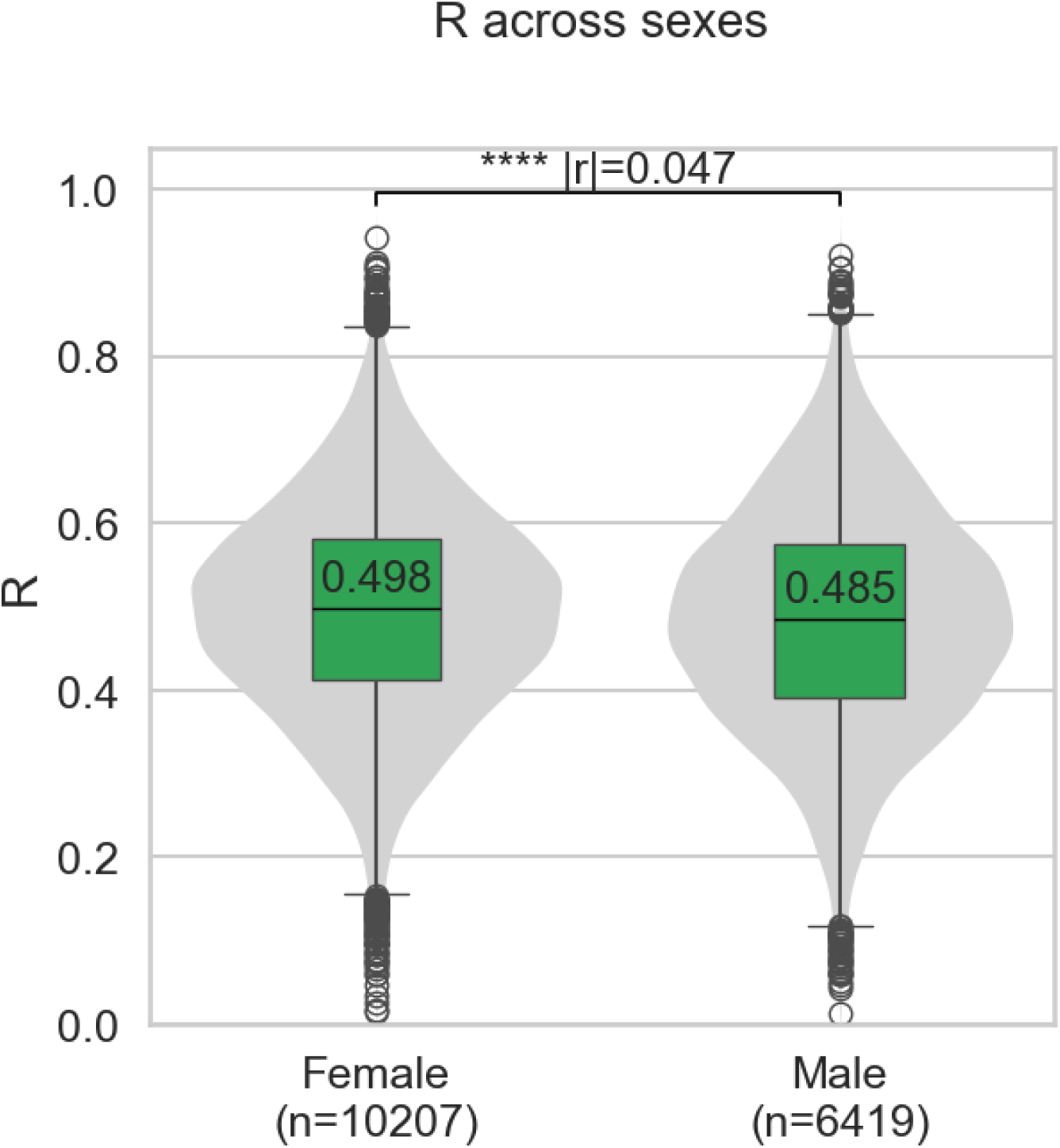

| Comparison | Statistic | p-value | $\epsilon^2$ / \|r\| | Median (Group 1) | Median (Group 2) |
| --- | --- | --- | --- | --- | --- |
| Overall | H = 25.704 | 3.981e-07 | 0.001 |  |  |
| Pairwise |  |  |  |  |  |
| Male vs Female | U = 31231816.0 | 3.981e-07 | 0.047 | 0.485 | 0.498 |

### S9. Group differences in R - pairwise comparison between chronotypes

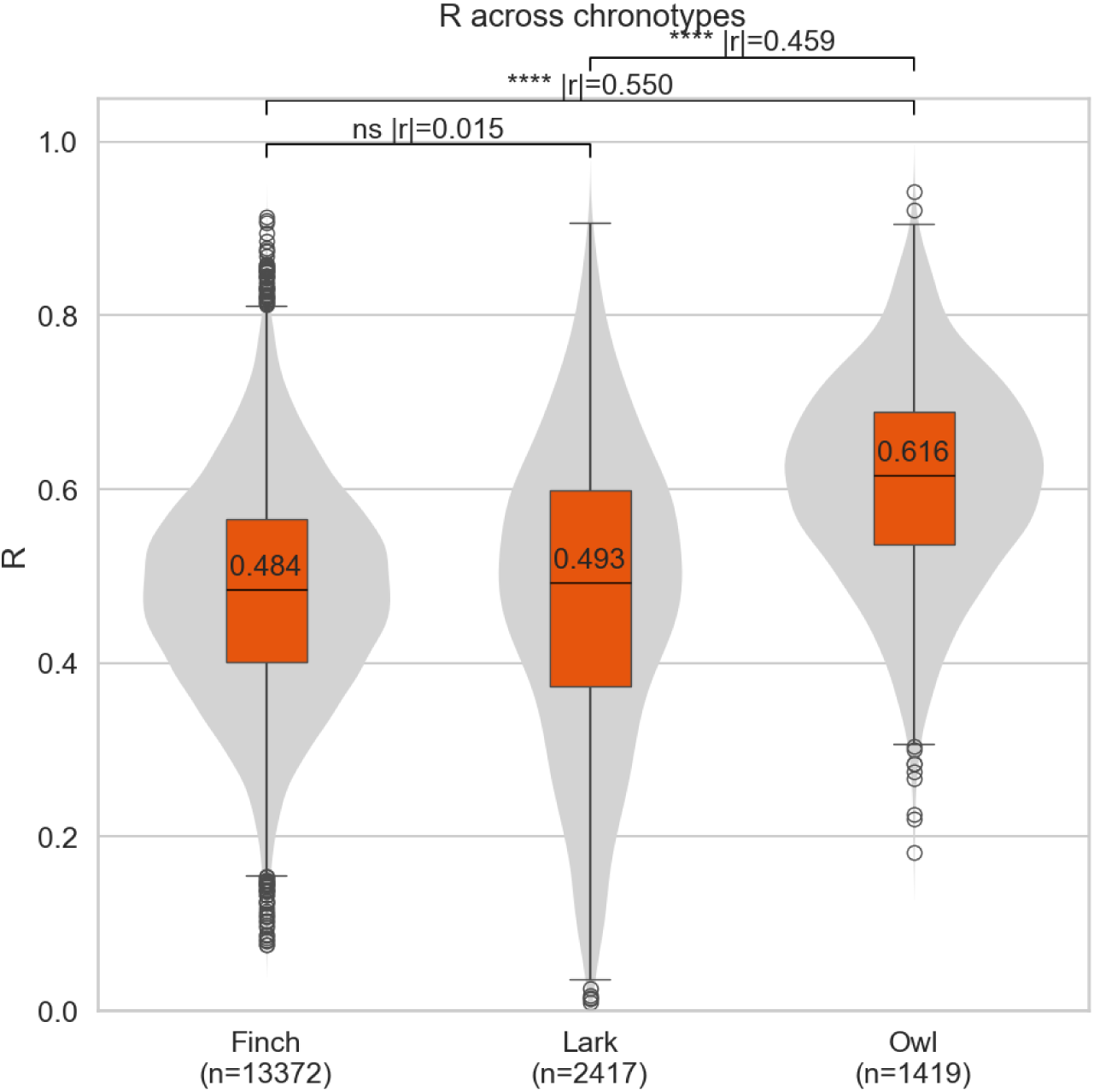

| Comparison | Statistic | p-value (FDR corrected) | $\epsilon^2$ / \|r\| | Median (Group 1) | Median (Group 2) |
| --- | --- | --- | --- | --- | --- |
| Overall | H = 1127.201 |  | 0.065 |  |  |
| Pairwise |  |  |  |  |  |
| Finch vs Lark | U = 15 914 464.0 | 2.34e-01 | 0.015 | 0.484 | 0.493 |
| Finch vs Owl | U = 4 264 702.0 | 4.07e-255 | 0.550 | 0.484 | 0.616 |
| Lark vs Owl | U = 927 088.0 | 6.60e-125 | 0.459 | 0.493 | 0.616 |

### S10. GEE Model equation for GPA vs R

We used mean-centered *R̅* values (*R̅*_cent_ = *R̅* – *R̅*_mean_), so *R̅*_cent_ = 0 represents “average distinctness” in our sample. Due to this procedure, the model intercept is “expected GPA at average regularity.”, instead of “expected GPA at completely spread activity (*R̅* = 0)”, which does not exist in real circadian behavior. Centering makes the intercept meaningful biologically. Additionally, centering orthogonalizes the linear and quadratic terms, leading to a reduction of multicollinearity between *R̅* and *R̅*^2^.

We modeled term GPA as a function of *R̅*_cent_ using Generalized Estimating Equations (GEE) with a Gaussian family, identity link, and an exchangeable working correlation to account for repeated semesters within students:

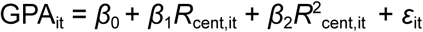

where GPA – student’s grade, *R̅*_cent_ – mean centered student’s distinctness, *i* – the student ID (grouping variable), *t* – the semester (observation within that student), *β*_0_,*β*_1_,*β*_2_ – model coefficients: the *average* intercept, linear, and quadratic term, *ε*_it_ – the residual term.

## Results

| Parameter | Coefficient | Std. Error | p-value | 95% CI (Lower) | 95% CI (Upper) |
| --- | --- | --- | --- | --- | --- |
| Intercept | 3.2554 | 0.009 | <0.001 | 3.238 | 3.272 |
| $\bar{R}_c$ | -0.0650 | 0.048 | 0.174 | -0.159 | 0.029 |
| $\bar{R}_c^2$ | -1.4564 | 0.245 | <0.001 | -1.936 | -0.977 |

### Model Information (GEE)

Observations: 17,376, Clusters (students): 9,014, Cluster size (min–max): 1–4, Mean cluster size: 1.9, Family: Gaussian, Link function: Identity, Working correlation: Exchangeable, Covariance type: Robust

### Distribution Diagnostics

Skewness: –1.500, Kurtosis: 2.878, Centered skewness: –0.628, Centered kurtosis: 7.828

**GEE model and Seasonal differences - season does not impact quadratic relationship with R**

|  | Coefficient | Std. Error | p-value | 95% CI (Lower) | 95% CI (Upper) |
| --- | --- | --- | --- | --- | --- |
| Intercept | 3.2560 | 0.009 | 0.000 | 3.239 | 3.273 |
| $\bar{R}_c \times \text{Season[Fall]}$ | -0.1083 | 0.060 | 0.070 | -0.225 | 0.009 |
| $\bar{R}_c \times \text{Season[Spring]}$ | -0.0375 | 0.064 | 0.557 | -0.163 | 0.088 |
| $\bar{R}_c^2 \times \text{Season[Fall]}$ | **-1.1034** | **0.279** | **0.000** | **-1.649** | **-0.557** |
| $\bar{R}_c^2 \times \text{Season[Spring]}$ | **-1.8325** | **0.330** | **0.000** | **-2.479** | **-1.186** |

**GEE model and Sex differences - sex does not impact quadratic relationship with R**

|  | Coefficient | Std. Error | p-value | 95% CI (Lower) | 95% CI (Upper) |
| --- | --- | --- | --- | --- | --- |
| Intercept | 3.2715 | 0.008 | 0.000 | 3.255 | 3.288 |
| $R_c \times \text{Sex}[\text{Male}]$ | -0.0291 | 0.072 | 0.685 | -0.170 | 0.112 |
| $R_c \times \text{Sex}[\text{Female}]$ | -0.0782 | 0.058 | 0.176 | -0.191 | 0.035 |
| <u>$R_c^2 \times \text{Sex}[\text{Male}]$</u> | <u>-1.9047</u> | <u>0.317</u> | <u>0.000</u> | <u>-2.526</u> | <u>-1.284</u> |
| <u>$R_c^2 \times \text{Sex}[\text{Female}]$</u> | <u>-0.7514</u> | <u>0.300</u> | <u>0.012</u> | <u>-1.339</u> | <u>-0.164</u> |

### S11. QIC results for GEE models for R-GPA relationship within each chronotype group

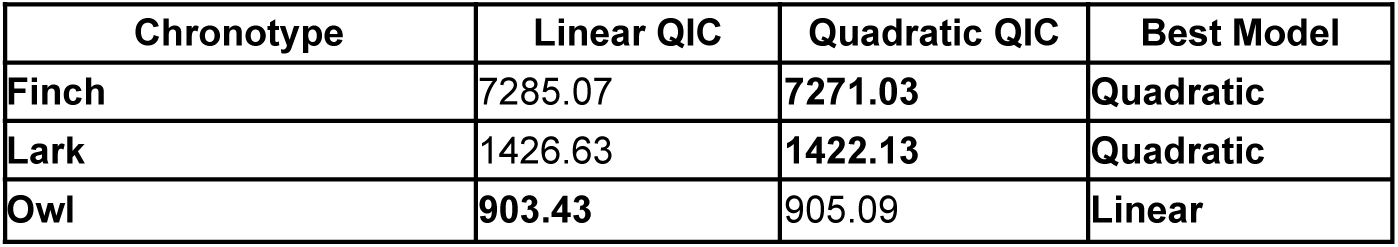

| Chronotype | Linear QIC | Quadratic QIC | Best Model |
| --- | --- | --- | --- |
| Finch | 7285.07 | **7271.03** | Quadratic |
| Lark | 1426.63 | **1422.13** | Quadratic |
| Owl | **903.43** | 905.09 | Linear |

### S12. GEE models for R-GPA relationship - different relationships for each chronotype

Lark

S12. GEE models for R-GPA relationship - different relationships for each chronotype Lark
|  | Coefficient | Std. Error | p-value | 95% CI (Lower) | 95% CI (Upper) |
| --- | --- | --- | --- | --- | --- |
| Intercept | 3.3511 | 0.21 | 0.000 | 3.310 | 3.392 |
| <u>$R_c$</u> | <u>0.3680</u> | <u>0.98</u> | <u>0.000</u> | <u>0.176</u> | <u>0.560</u> |
| <u>$R_c^2$</u> | <u>-1.0643</u> | <u>0.442</u> | <u>0.016</u> | <u>-1.931</u> | <u>-0.198</u> |

Finch

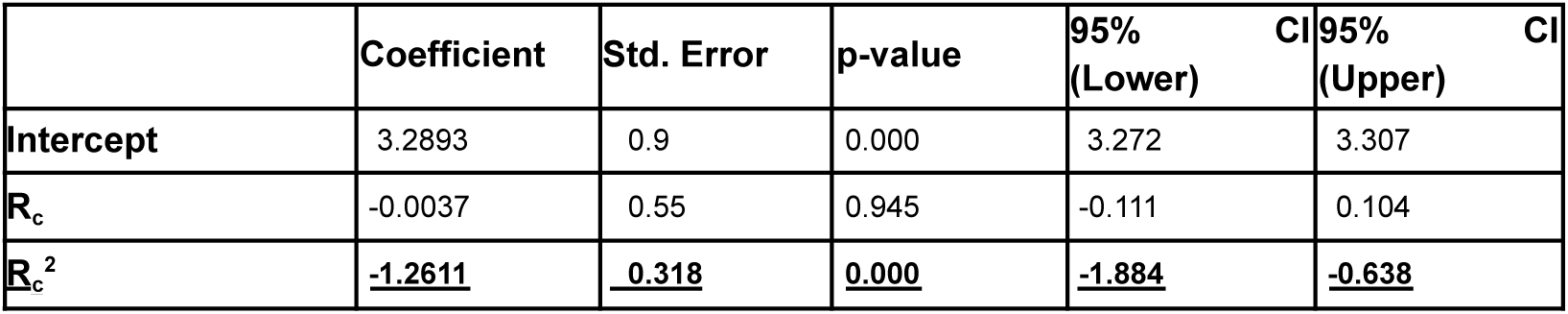

Owls

|  | Coefficient | Std. Error | p-value | 95% CI (Lower) | 95% CI (Upper) |
| --- | --- | --- | --- | --- | --- |
| Intercept | 3.0655 | 0.023 | 0.000 | 3.021 | 3.110 |
| $R_c$ | -0.3328 | 0.176 | 0.059 | -0.678 | 0.013 |

### S13. GEE model for the relationship between R and SJL

We modeled SJL as a function of *R̅*_cent_ using Generalized Estimating Equations (GEE) with a Gaussian family, identity link, and an exchangeable working correlation to account for repeated semesters within students:

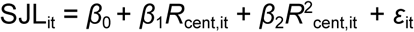

where SJL – student’s social jetlag, *R̅*_cent_ – mean centered student’s distinctness, *i* – the student ID (grouping variable), *t* – the semester (observation within that student), *β*_0_,*β*_1_,*β*_2_ – model coefficients: the *average* intercept, linear, and quadratic term, *ε*_it_ – the residual term.

| Parameter | Coefficient | Std. Error | p-value | 95%<br>(Lower) | 95%<br>(Upper) |
| --- | --- | --- | --- | --- | --- |
| Intercept | 0.4278 | 0.008 | 0.000 | 0.411 | 0.444 |
| $R_c$ | <u>1.1337</u> | <u>0.053</u> | <u>0.000</u> | <u>1.029</u> | <u>1.238</u> |
| $R_c^2$ | <u>-0.8474</u> | <u>0.270</u> | <u>0.002</u> | <u>-1.376</u> | <u>-0.319</u> |

### S14. QIC results for GEE models for R-SJL relationship within each chronotype group

| Chronotype | Linear QIC | Quadratic QIC | Best Model |
| --- | --- | --- | --- |
| Lark | 790.15 | 790.82 | Linear/Quadratic |
| Finch | 6064.39 | 6065.61 | Linear |
| Owl | 872.58 | 865.06 | Quadratic |

### S15. GEE models for R-SJL relationship - different relationships for each chronotype

**Lark**

S15. GEE models for R-SJL relationship - different relationships for each chronotype Lark
|  | Coefficient | Std. Error | p-value | 95%<br>(Lower) | 95%<br>(Upper) |
| --- | --- | --- | --- | --- | --- |
| Intercept | -0.3200 | 0.016 | 0.000 | -0.352 | -0.288 |
| $R_c$ | 0.1026 | 0.077 | 0.184 | -0.049 | 0.254 |
| $R_c^2$ | <u>-0.7143</u> | <u>0.347</u> | <u>0.040</u> | <u>-1.395</u> | <u>-0.034</u> |

**Finch**

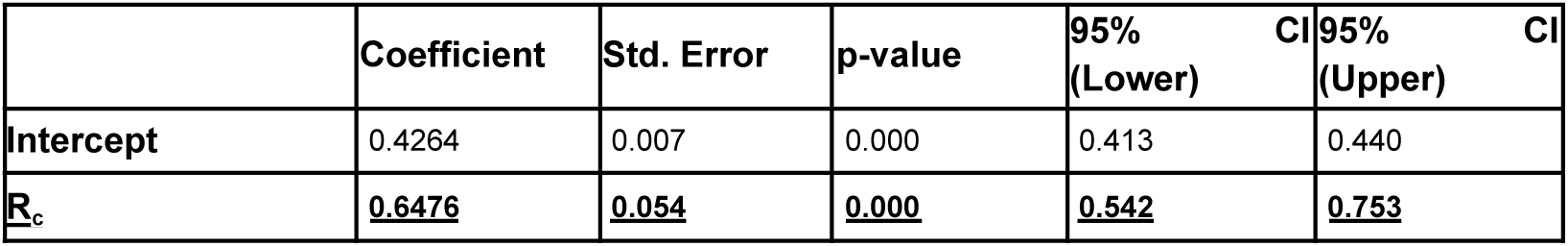

|  | Coefficient | Std. Error | p-value | 95%<br>(Lower) | 95%<br>(Upper) |
| --- | --- | --- | --- | --- | --- |
| Intercept | 0.4264 | 0.007 | 0.000 | 0.413 | 0.440 |
| $R_c$ | <u>0.6476</u> | <u>0.054</u> | <u>0.000</u> | <u>0.542</u> | <u>0.753</u> |

**Owl**

| Parameter | Coefficient | Std. Error | p-value | 95%<br>(Lower) | 95%<br>(Upper) |
| --- | --- | --- | --- | --- | --- |
| Intercept | 1.4113 | 0.026 | 0.000 | 1.360 | 1.463 |
| $R_c$ | <u>0.5219</u> | <u>0.191</u> | <u>0.006</u> | <u>0.148</u> | <u>0.896</u> |
| $R_c^2$ | <u>4.4343</u> | <u>1.035</u> | <u>0.000</u> | <u>2.405</u> | <u>6.464</u> |

### S15. GEE model for R-SJL-GPA relationship

We modeled term GPA as a function of *R̅* and SJL using Generalized Estimating Equations (GEE) with a Gaussian family, identity link, and an exchangeable working correlation to account for repeated semesters within students:

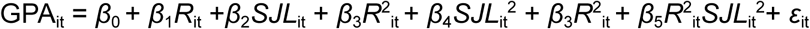

where GPA – student’s grade, *R̅* – distinctness, SJL – student’s social jetlag, *i* – the student ID (grouping variable), *t* – the semester (observation within that student), *β*_0_,*β*_1_,*β*_2,_*β*_3_,*β*_4_ – model coefficients: the *average* intercept, linear, and quadratic term, *ε*_it_ – the residual term.

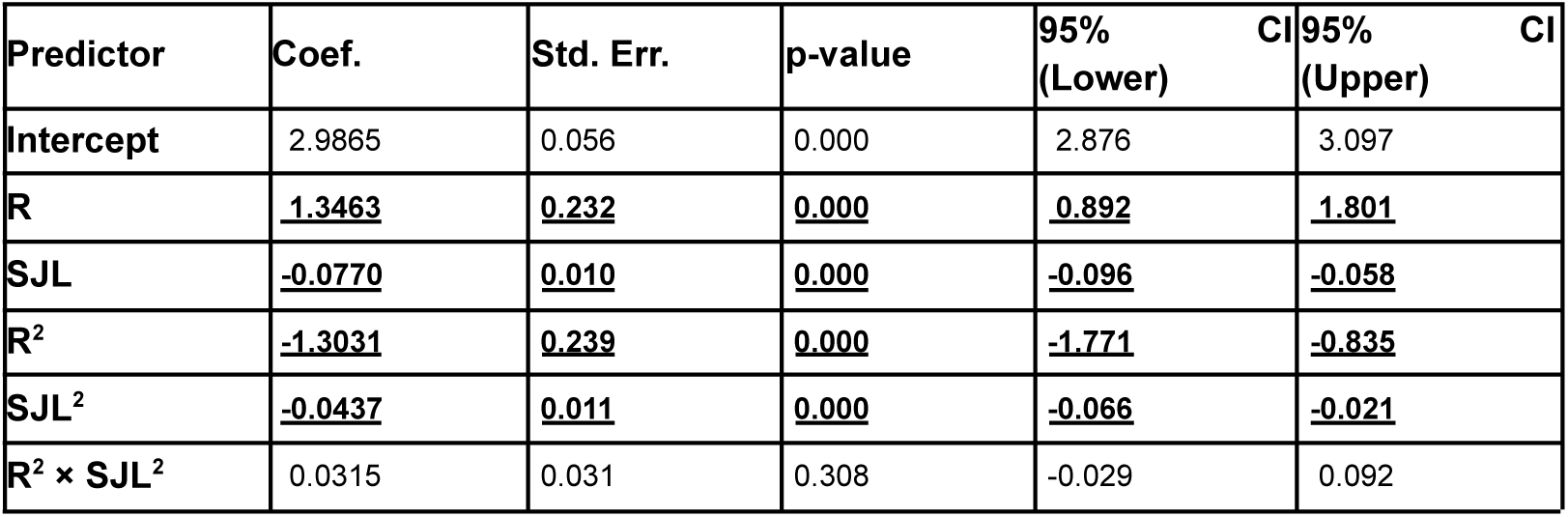

| Predictor | Coef. | Std. Err. | p-value | 95%<br>(Lower) | 95%<br>(Upper) |
| --- | --- | --- | --- | --- | --- |
| Intercept | 2.9865 | 0.056 | 0.000 | 2.876 | 3.097 |
| R | <u>1.3463</u> | <u>0.232</u> | <u>0.000</u> | <u>0.892</u> | <u>1.801</u> |
| SJL | <u>-0.0770</u> | <u>0.010</u> | <u>0.000</u> | <u>-0.096</u> | <u>-0.058</u> |
| $R^2$ | <u>-1.3031</u> | <u>0.239</u> | <u>0.000</u> | <u>-1.771</u> | <u>-0.835</u> |
| $SJL^2$ | <u>-0.0437</u> | <u>0.011</u> | <u>0.000</u> | <u>-0.066</u> | <u>-0.021</u> |
| $R^2 \times SJL^2$ | 0.0315 | 0.031 | 0.308 | -0.029 | 0.092 |

### S16. GEE model for R-SJL-GPA relationship - within chronotype groups

**Larks**

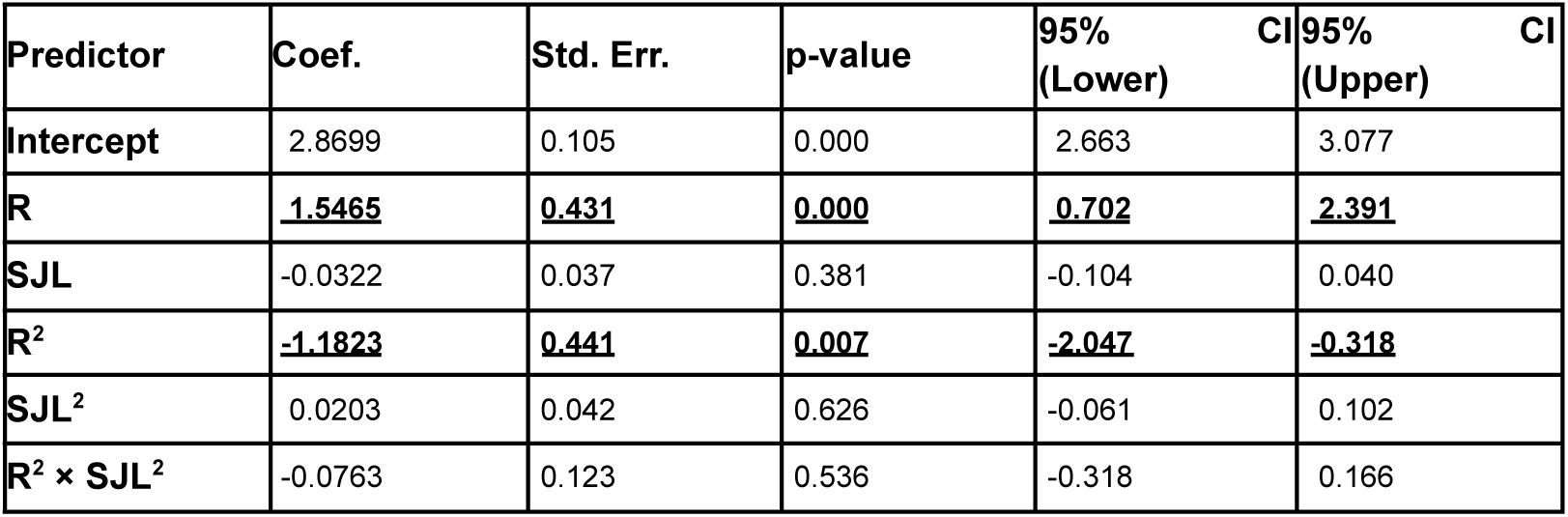

**Finches**

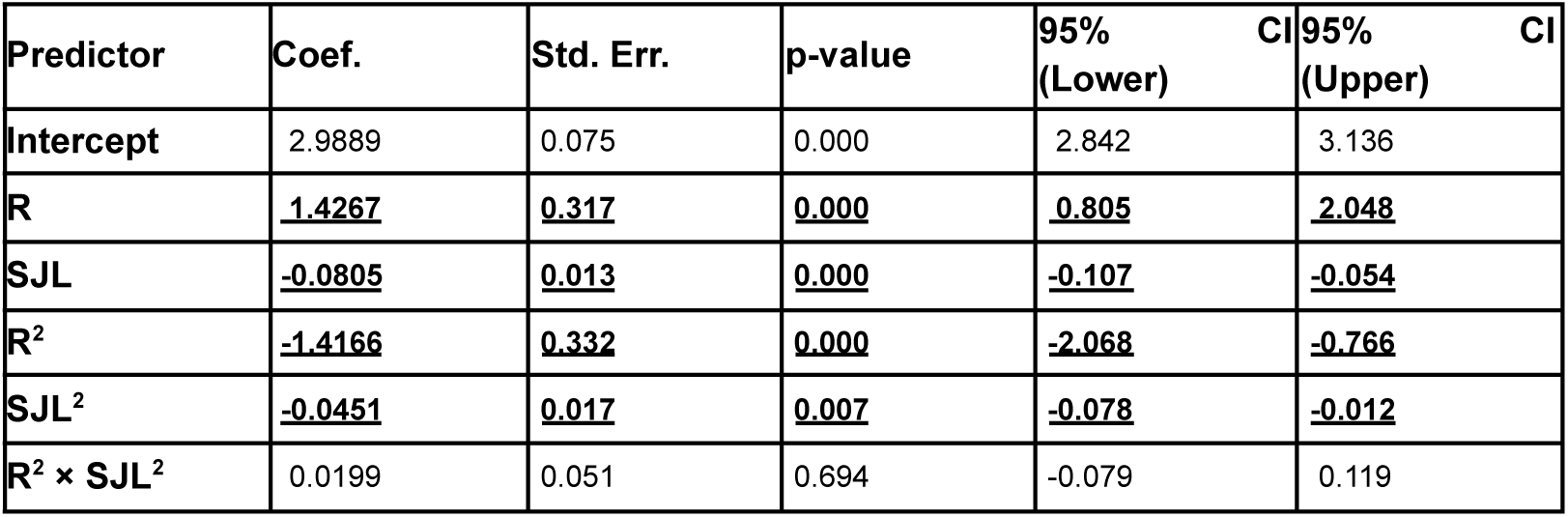

**Owls**

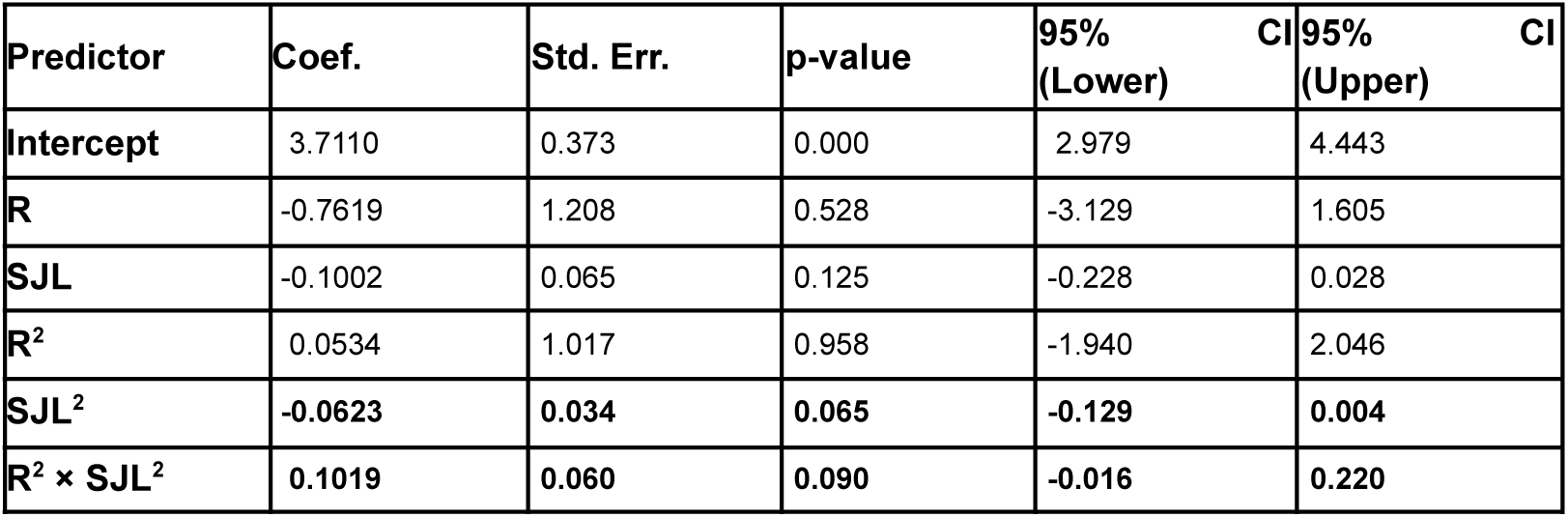

### S17. Goodness-of-fit estimates (pseudo-R²) for chronotype-specific GEE models for R-SJL-GPA relationship

S17. Goodness-of-fit estimates (pseudo-R<sup>2</sup>) for chronotype-specific GEE models for R-SJL-GPA relationship.
| Chronotype | Zhang<br>pseudo-R<sup>2</sup> | Cluster-mean<br>correlation<br>pseudo-R<sup>2</sup> |
| --- | --- | --- |
| Finch | 0.0165 | 0.0338 |
| Lark | 0.0133 | 0.0158 |
| Owl | 0.0316 | 0.0392 |

